# Assembly and cell-free expression of a partial genome for the synthetic cell

**DOI:** 10.1101/2025.10.16.682769

**Authors:** Céline Cleij, Laura Sierra Heras, Ellen Zwiers, Etienne Bancal, Alexandre Stella, Pascale Daran-Lapujade, Christophe Danelon

## Abstract

The de novo design and assembly of a DNA genome represents an important milestone toward the construction of a minimal synthetic cell. The genome must contain all the instructions to enable primary cellular functions, probably comprising over 150 genes with a total size over 200 kb. Here, we designed and built a partial synthetic genome that satisfies the requirements for its expression in PURE system—a minimal transcription-translation machinery reconstituted from purified elements—and its replication by the protein-primed φ29 DNA replication system. The partial minimal synthetic genome (MSG) has a size of 41 kb, was assembled in yeast from 14 fragments, and harbors genes for phospholipid biosynthesis, DNA replication, and cell division. Yeast marker fragments were added between coding fragments to facilitate screening of correct assemblies and a BAC backbone was included to enable transfer from yeast to *E. coli* for amplification. Synthesis of all MSG-encoded proteins in PURE system was confirmed by liquid chromatography-mass spectrometry and fluorescence measurements. Moreover, we demonstrate successful compartmentalization and expression of the MSG in liposomes, as well as full-length replication of the linearized MSG by the φ29 DNA replication machinery. This work provides proof-of-concept for the bottom-up assembly and cell-free expression of a functional genome for a minimal synthetic cell.

## Introduction

The bottom-up construction of synthetic cells requires the design and assembly of a DNA genome encoding all cellular functions. The exact gene content, organization, and DNA size that would support life in its simplest form are currently unknown. Estimates range from the theoretical approximations of 150–400 protein-coding genes ^1–9^ to an upper limit given by the 531-kb genome of the JCVI-syn 3.0 cell, containing 473 genes (of which 438 are protein-coding), which has been designed through genome reduction of *Mycoplasma mycoides* ^10^. In this latter study, a minimal bacterial genome was constructed from shorter DNA fragments through a combination of in vitro assembly followed by homologous recombination in yeast. This shows the feasibility of stepwise assembly of genome-sized DNA. However, the design of this minimal genome was based on the native *M. mycoides*, largely maintaining the original sequence and gene order.

In contrast, we envision a synthetic cell genome comprising a set of genes and regulatory elements (native or modified) derived from various organisms, or even entirely new to nature ^11^. Modules are selected based on their apparent simplicity, prioritizing mechanisms that achieve a given function with the fewest necessary genes. An additional selection criterion is the compatibility with the core biosynthesis machinery. In this context, Protein Synthesis Using Recombinant Elements (PURE) system ^12,13^ is particularly relevant because of its reconstituted nature, well-defined composition, and minimal number of proteins and cofactors to support gene expression. In the commercial version of PURE system, transcription is controlled by the T7 RNA polymerase, while translation relies on *E. coli* ribosomes and factors.

Important milestones toward the completely de novo design and assembly of a chromosome for a synthetic cell are the synthesis of a 15-kb DNA construct containing all 21 genes that encode the proteins of the *E. coli* 30S ribosomal subunit ^14^ and the construction of a 30-cistron translation factor module, named pTFM1, encoding 30 of the 31 translation factors of PURE system, with only EF-Tu missing ^15^. Although successful and relatively cheap, the assembly of pTFM1 by BioBrick cloning was labor-intensive, posing a challenge for scaling up the number of genes.

A suitable method for the construction of a minimal synthetic genome should thus enable the assembly of >100-kb DNA from multiple fragments, and preferably allow for the modular, one-pot, assembly of transcription cassettes to ease testing a variety of genome designs. Despite recent successes to develop in vitro DNA assembly methods ^15–17^ (reviewed in ^18^), this approach is currently not compatible with the intended genome size and number of transcription units. The most well-established method for one-pot assembly of multiple fragments into large constructs is in vivo assembly in the yeast *Saccharomyces cerevisiae*. This homology-based assembly approach was used for the construction of several genomes in the JCVI *Mycoplasma* project ^10,19,20^. It is capable of one-pot assembly of 44 fragments ^21^, and is suitable for the assembly of chromosomes up to 1.66 Mb ^22^ (reviewed in ^23^). By including homologous overhangs in the assembly fragments, the homology-directed repair (HDR) machinery of *S. cerevisiae* can be employed to stitch linear DNA fragments upon transformation. Moreover, the use of synthetic homology regions (SHRs) enables modularity of the assembly design ^24^. Yet, the capability of *S. cerevisiae* to correctly assemble DNA parts containing heterologous sequences with multiple repeats, such as T7 regulatory sequences and canonical prokaryotic RBSs, remains unexplored.

In this study, we establish yeast as an assembly platform to construct minimal genomes for PURE-based synthetic cells. We designed mock cassettes containing PURE regulatory sequences at both ends, and included SHRs through PCR. After transformation of yeast with 20 DNA fragments, correct assemblies were identified using selection markers and verified by long-read DNA sequencing, which provided insights into the role of repeats in unintended recombination events. The 67-kb synthetic chromosome (SynChr) was isolated from yeast and directly assayed for expression in PURE system. Based on these results, we constructed a partial minimal synthetic genome, MSG1, harboring 15 genes involved in basic cellular functions inspired from *E. coli* and the bacteriophage φ29: phospholipid synthesis, DNA replication, and cell division. The SynChr included a BAC backbone to enable shuttling to *E. coli* for amplification and isolation with high DNA yield. Purified MSG1 was expressed in PURE system and the synthesis of all encoded proteins was verified by mass spectrometry and fluorescence. We demonstrated that MSG1 could be expressed in liposomes, linearized, and replicated by the protein-primed replication mechanism of phage φ29, completing all steps of the central dogma.

## Results

### Design of a mock synthetic chromosome with repeated sequences

We have explored homologous recombination in the yeast *Saccharomyces cerevisiae* as a method for the assembly of SynChrs from multiple DNA fragments. The compatibility of this approach with the constraints imposed by the utilization of PURE system for cell-free gene expression was first tested by designing a mock SynChr, called MSG0.1, harboring the following features: i) the presence of multiple transcription units for expression in PURE system, ii) the ability to be assembled and maintained in yeast, and iii) the possibility for easy screening of yeast clones with correctly assembled SynChrs. In addition, MSG0.1 was equipped with an *E. coli* replication origin, an optional feature in case amplification of the SynChr would be required.

The designed MSG0.1 includes ten transcriptional units. Each cassette starts with the same 139-bp sequence containing a T7 promoter and an *E. coli* RBS, and ends with the same 119-bp sequence containing a T7 terminator, hereafter called ‘PURE repeats’ (Figure 1). One of these cassettes contains the reporter gene *yfp* for characterization of protein synthesis in PURE system. The other nine cassettes were designed as mock transcription units with standardized DNA parts. They do not lead to expressed proteins in PURE and were designed to be non-coding in both *S. cerevisiae* and *E. coli* to avoid expression of potentially toxic proteins. They consist of 5-kb genomic sequences from the plant *Arabidopsis thaliana* flanked by PURE repeats. Expression of plant DNA is not expected in yeast ^25^ and, while transcription and translation may not totally be eliminated in *E. coli*, this design should keep it to a minimum ^26–31^. The *A. thaliana* DNA fragments were checked for the absence of T7 promoters to prevent transcription in PURE system.

**Figure 1:**
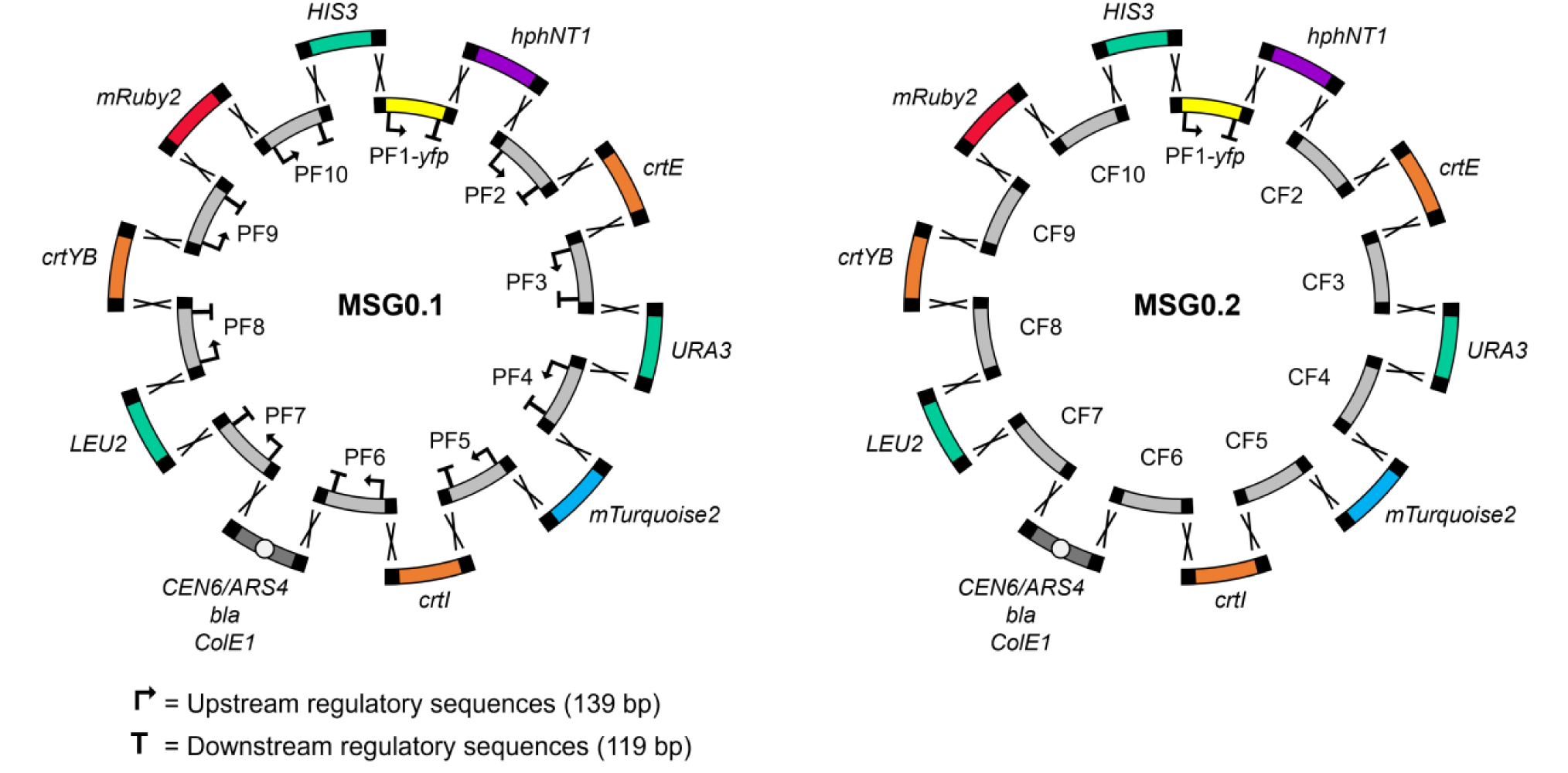
Design of a synthetic chromosome for expression in PURE system. Design of MSG0.1 (left) and MSG0.2 (right), differing only by the absence of PURE repeats in CF2–CF10. PF, PURE fragment with repeats; CF, control fragment without repeats.

To simplify the screening pipeline for identification of yeast strains harboring correctly assembled SynChrs, a series of nine genetic selection and screening markers were included as DNA parts interspacing the PURE transcription cassettes (Figure 1): *hphNT1* conferring resistance to hygromycin, *crtYB*, *crtE,* and *crtI* leading to β-carotene biosynthesis identifiable by the orange coloring of colonies in the presence of all three genes, *mTurqoise2* and *mRuby2* encoding fluorescent proteins, and three auxotrophic markers *URA3*, *LEU2,* and *HIS3* enabling growth in the absence of uracil, leucine, and histidine, respectively.

For replication and segregation of MSG0.1 in yeast, a centromeric origin and an autonomously replicating sequence were added to the design as a single fragment (*CEN6/ARS4*). We therefore expected MSG0.1 to be present in a single copy per cell on average. Alternatively, we explored a design incorporating a 2µ replication origin (MSG0.1^2µ^) to increase the copy number and to allow higher yields during extraction. Finally, a bacterial ColE1-type replication origin and the *bla* antibiotic resistance gene (conferring resistance to ampicillin) were added on the same DNA fragment as the yeast origin of replication, giving the possibility to maintain and amplify the SynChr in *E. coli*. A control SynChr (MSG0.2) was also designed, differing only in the absence of PURE repeats, with a total size of 66 kb (Figure 1). Similarly, a control SynChr with 2µ origin was designed (MSG0.2^2µ^). All designed SynChr maps are available in Supplementary Data 5.

### Assembly, extraction from yeast, and cell-free expression of the mock synthetic chromosome

We established a pipeline to screen, prior to long-read DNA sequencing, for yeast strains harboring correctly assembled SynChrs (Figure 2). The screening pipeline was designed to exclude PCR verification for two reasons: the inherent risk of false negative results, and in case of low assembly efficiency, colony screening by PCR would be excessively labor-intensive.

**Figure 2:**
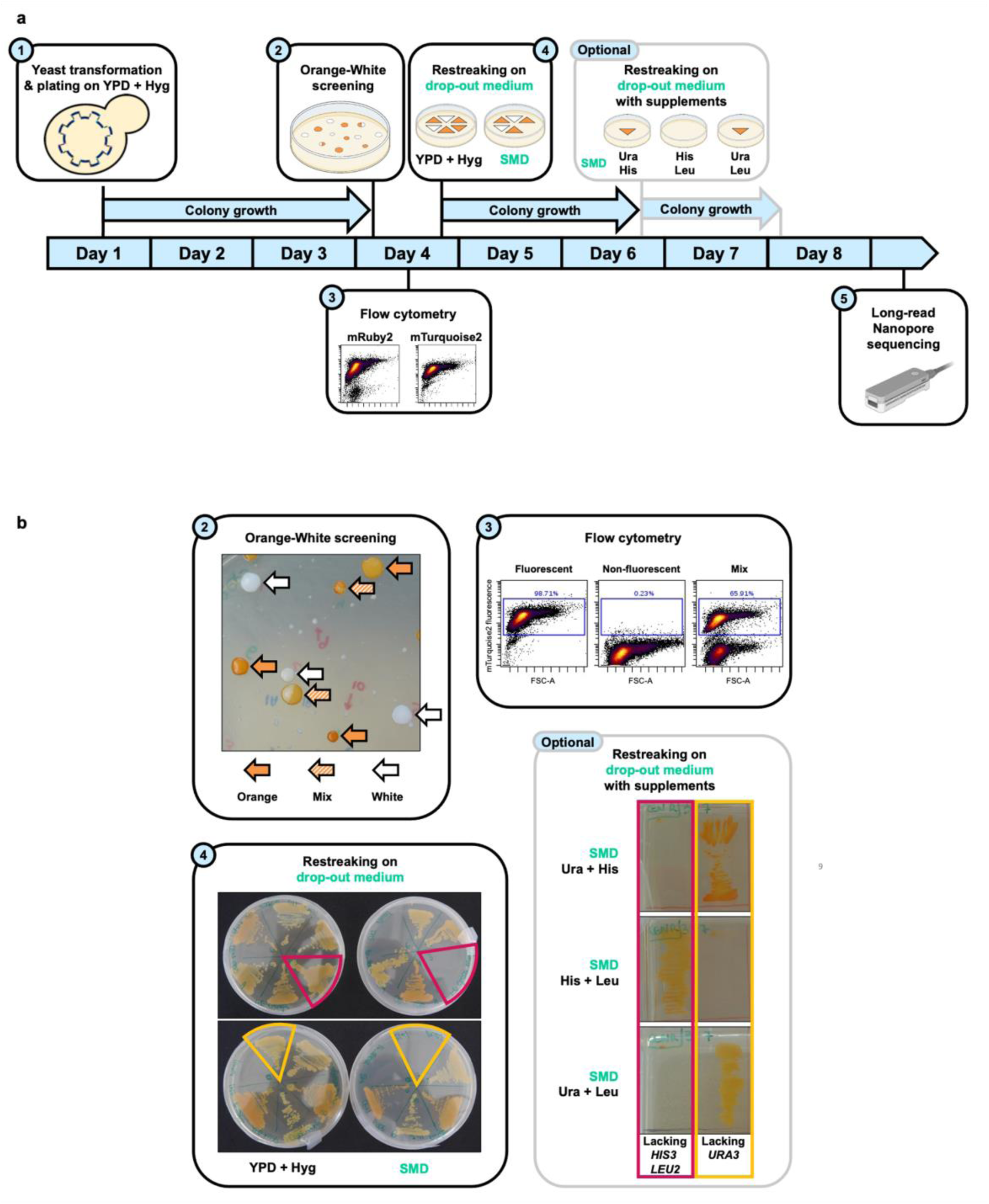
Screening pipeline to select clones with correct assemblies. **a)** Step 1: Transformation of yeast with DNA fragments and plating on selective agar plates containing YPD and hygromycin. Visible colonies are formed within three days. Step 2: Orange-White screening by visual inspection, based on β-carotene production. Step 3: mRuby2 and mTurquoise2 detection by flow cytometry. Step 4: Detection of auxotrophies by restreaking on YPD + Hyg and SMD. Growth is visible within two days. Optional step: in case of absence of growth on SMD, specific auxotrophies can be identified by restreaking on various drop-out media followed by growth for two days. Step 5: Positively screened strains were selected for long-read Nanopore sequencing. **b)** Panels with classification examples for steps 2, 3, 4 and optional screening step.

After transformation and assembly in yeast, plating on YPD with hygromycin selected for strains with SynChrs carrying the *hphNT1* fragment (Figure 2, step 1). Visual inspection of the colonies after three days further revealed the presence of all three carotenoid genes (Figure 2, step 2), as their simultaneous presence leads to an orange coloring of the colonies ^32^. Fluorescence analysis by flow cytometry identified clones expressing the mRuby2 and mTurquoise2 proteins (Figure 2, step 3). Finally, plating on minimal synthetic medium selected for colonies harboring all three auxotrophic markers (Figure 2, step 4). For additional information, plating on media lacking single nutrients could be used to identify the absence of specific auxotrophic markers. The entire screening pipeline was designed to test hundreds of clones within six days for the presence or absence of the nine markers. This would allow identification of correct assemblies with occurrence frequencies as low as 0.3–1%. Positive clones were sequenced using Nanopore technology (Figure 2, step 5).

Three independent yeast transformations were conducted with the 20 assembly fragments for both the MSG0.1 and MSG0.2. The total number of colonies over three transformations was considered rather than individual transformations to report the assembly efficiency, as this mitigates expected day-to-day variation between independent transformations. Detailed results for the different screening steps are reported in Supplementary Note 1 and Supplementary Figures 1 and 2. We found that correct assembly of MSG0.1 may be possible, albeit with a lower efficiency than that of control MSG0.2 (9% vs. 86% of total colony count in step 1 passed screening steps 1–4), indicating that PURE repeats cause undesired recombination events. Moreover, preliminary data revealed a low assembly efficiency of both MSG0.1^2µ^ and MSG0.2^2µ^, regardless of the presence of repeats (Supplementary Note 1 and Supplementary Figure 3). Therefore, we did not further investigate SynChr assembly with 2µ replication origins.

The five strains identified by the screening pipeline as harboring MSG0.1 with all expected marker fragments were sequenced by long-read Nanopore sequencing (Supplementary Data 1.2). Four out of the five strains showed correct SynChr configurations. In the remaining strain, recombination events occurred between PURE repeats as revealed by the raw reads, but no consensus sequence could be determined. Details of the sequencing results can be found in Supplementary Note 2 and Supplementary Figures 4–6. Overall, Nanopore sequencing confirmed that *S. cerevisiae* is capable of successfully assembling SynChrs from at least ten fragments with internal sequence repeats, albeit with a relatively low efficiency (ca. 8%). The established screening pipeline allowed us to select positive clones with the desired configuration of SynChrs, alleviating the need for extensive sequencing or colony PCR.

Notably, for subsequent isolation, it is essential that SynChrs do not undergo recombination during propagation in yeast. MSG0.1 and MSG0.2 are stably maintained for at least 32 generations and the presence of PURE repeats does not hamper stability (Supplementary Note 3 and Supplementary Figure 7).

Next, we investigated the ability of MSG0.1 to serve as template for *yfp* expression in PURE system. Several iterations of protocol improvement were necessary to extract a sufficient amount of MSG0.1 from yeast and achieve detectable fluorescence levels of YFP. Starting from about 20 pM of MSG0.1 template, clear production of YFP was measured after 16 h of incubation compared to the negative control containing water instead of DNA (Supplementary Note 4, Supplementary Figure 8c). Yet, protein synthesis yield was low, indicating that further optimization is required to increase the concentration and purity of SynChrs for efficient expression in PURE system.

### Design of a partial synthetic minimal genome

Having established a yeast-based assembly protocol for a mock SynChr that can be expressed in PURE system, we set out to build a partial minimal genome (MSG1) encoding three cellular modules, namely membrane growth, cell division, and DNA replication (Figure 3). These modules were previously characterized when individually expressed in PURE system. The membrane growth module consisted of genes encoding enzymes of the *E. coli* Kennedy pathway for phospholipid synthesis: *plsB*, *plsC*, *cdsA*, *pssA*, *psd*, *pgsA* and *pgpA* ^33^. Genes involved in two bacterial cell division processes, both from *E. coli*, were included: *minD* and *minE* encoding Min system proteins that can localize the division machinery and deform liposomes ^34,35^, and *ftsZ* and *ftsA* encoding divisome proteins which together can constrict liposomes ^36,37^. FtsZ was fused to the fluorescent mVenus protein for easy visualization during in vitro experiments, and the *ftsZ-mVenus* and *ftsA* genes were organized into an operon. The DNA replication module consisted of the *p2* and *p3* genes encoding DNAP and TP of the protein-primed DNA replication system from bacteriophage φ29 ^38^. Expression of all genes for the synthetic cell was under the control of a T7 promoter (pT7), with the exception of the phospholipid synthesis genes *pgsA* and *pgpA* that were placed under the control of an SP6 promoter (pSP6) ^33^. SP6 RNAP can be added to PURE system and is expected to have orthogonal activity to T7 RNAP ^33^. The presence of SP6 and T7 promoters enables separate activation of the two branches of the phospholipid synthesis pathway. Additionally, MSG1 design included fluorescent markers for direct readout of protein synthesis in PURE system: *mCherry* controlled by pT7, and *eYFP* controlled by pSP6. The fusion protein FtsZ:mVenus could also be used as fluorescent readout under pT7 control. The chosen DNA replication machinery requires a linear DNA molecule with φ29 origins at both ends. Therefore, MSG1 included a fragment containing the two origins of replication with an internal PmeI restriction site for linearization. The presence of a PmeI site imposes an extra design constraint, i.e., the absence of additional PmeI recognition sites on the SynChr, which is the case for MSG1.

**Figure 3:**
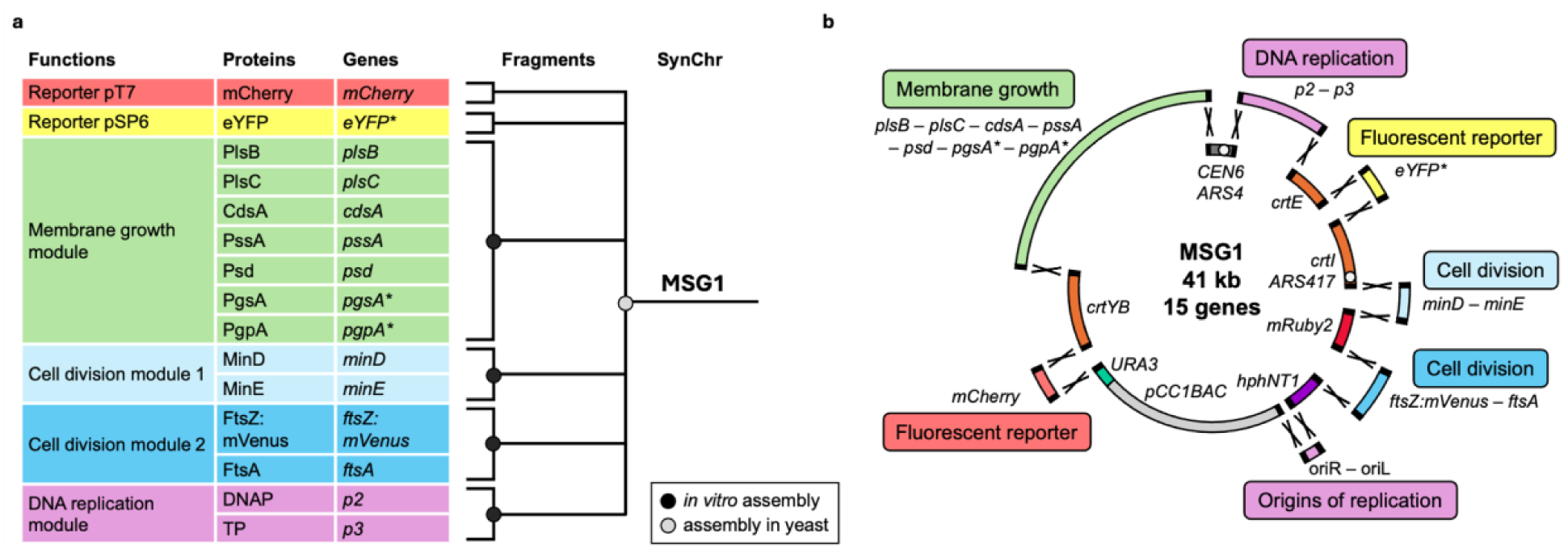
Design of a partial minimal synthetic genome. **a)** Genes and encoded proteins on MSG1. Genes involved in each cellular module were previously cloned together on a plasmid in vitro (Supplementary Data 2.3). Fragments were amplified from these plasmids by PCR and assembled in yeast to obtain MSG1. **b)** Design of MSG1 encoding 15 genes for expression in PURE system and with a total size of 41 kb. A marker for screening and selection in yeast or *E. coli* was introduced between each fragment harboring a synthetic cell module. Genes indicated with an asterisk are controlled by pSP6, all other genes by pT7.

The regulatory sequences (promoter, RBS, terminator) of most genes are similar and therefore repeated multiple times across MSG1, which is expected to cause unwanted recombination during assembly in yeast as shown with MSG0.1. To minimize the number of transformants that needed to be sequenced for identification of correct assemblies, yeast markers (auxotrophic, fluorescent, antibiotic resistance and chromogenic markers) were included between each minimal cell module according to the design of MSG0.1. To ensure maintenance in yeast, the designs included a *CEN6* centromere and an *ARS4* replication origin, along with one additional autonomously replicating sequences (*ARS417*), providing a replication origin every 30–40 kb ^39^. The *CEN6*/*ARS4* fragment additionally contained a landing pad with synthetic gRNA target sites ^40,41^ to allow for in vivo engineering by CRISPR/Cas9 in future studies. Due to the low concentration and purity obtained when isolating MSG0.1 from yeast, an amplification step in *E. coli* was envisioned. MSG0.1 and MSG0.2, which contained a high-copy number ColE1-like origin of replication, could not be stably maintained in *E. coli*. This was likely due to the incompatibility of the replication origin with the large SynChr size and to undesired recombination at the repeated regulatory sequences of MSG0.1. Large inserts can be stably maintained in bacterial artificial chromosomes (BACs), because their low copy number alleviates metabolic burden, limits intermolecular recombination, and provides increased tolerance for toxic sequences ^42,43^. Therefore, a pCC1BAC backbone (Epicentre) was included in the design of MSG1. Constructs with a pCC1BAC backbone are maintained at a single copy, but the optional activation of a second origin through *trfA* expression in *E. coli* strain EPI300—induced by L-arabinose—raises the copy number to approximately 10–20, enhancing DNA yield and purity upon isolation (Epicentre, ^44^).

MSG1 therefore comprised 14 assembly fragments: seven fragments containing minimal cell modules or reporters for expression in PURE system, which have previously been constructed by in vitro assembly (Supplementary Data 2.3), and seven marker fragments for propagation, screening and selection in yeast and *E. coli*. All fragments were flanked by 60-bp SHR sequences for assembly in yeast through homologous recombination. Final MSG1 was 41 kb in size and included 15 genes for expression in PURE system (Figure 3). The full content of the SynChr is described in Supplementary Data 4.1. Assembly fragments are listed in Supplementary Data 4.3 and a list of the encoded proteins for expression in PURE system is provided in Supplementary Data 4.4.

### Assembly in yeast and isolation of MSG1 at nanomolar concentrations

A single transformation was carried out using the 14 assembly fragments, which resulted in approximately 65 colonies on the transformation plate (Supplementary Figure 9). Out of the 35 orange colonies, 28 were subjected to the full screening and selection pipeline described above, resulting in identification of 24 colonies that contained all markers. After total DNA isolation from seven yeast strains and long-read sequencing, three strains were selected for further investigation, which harbored versions of MSG1 that were close to the design (MSG1.1, MSG1.2 and MSG1.3, Supplementary Data 1.2). MSG1.1 contained all fragments in the correct configuration, whereas MSG1.2 lacked a single gene (*pgpA*). MSG1.3 consisted of a mixture of correctly assembled SynChrs and some that were missing *mCherry* and the lipid synthesis genes. All three versions showed a relatively high occurrence of point mutations in the lipid synthesis segment due to the low-fidelity DNA polymerase used in this PCR (Supplementary Data 4.3 and 4.5).

Total DNA was isolated from yeast strains carrying MSG1 and used to transform *E. coli* strain EPI300 for amplification (Figure 4a). Transformation was successful for all three MSG1 variants. Single colonies from the transformation plates were checked for the presence of pCC1BAC, *mRuby2* and *CEN6*/*ARS4* fragments by colony PCR, and positive colonies were cultured without high-copy number induction. DNA was isolated from *E. coli* and sequenced, revealing a mixture of complete and incomplete SynChrs that were missing sequences between repeats. MSG1 may have been unstable during transformation of *E. coli* or subsequent recovery (as also observed in ^15^), leading to colonies on the transformation plate consisting of a mixed population of cells with different SynChr configurations. Therefore, cells from the transformation plates were streaked to obtain single colonies that were checked by PCR and used to inoculate overnight cultures. Since MSG1 is assumed to be stable during colony growth after streaking ^15^, these single colonies were expected to contain either recombined or full synthetic genomes, but not a mixture. This workflow ensured that cultures inoculated from a resuspended colony, which was confirmed as positive by colony PCR, contained a single configuration of MSG1. Sequencing confirmed single SynChr configurations.

**Figure 4:**
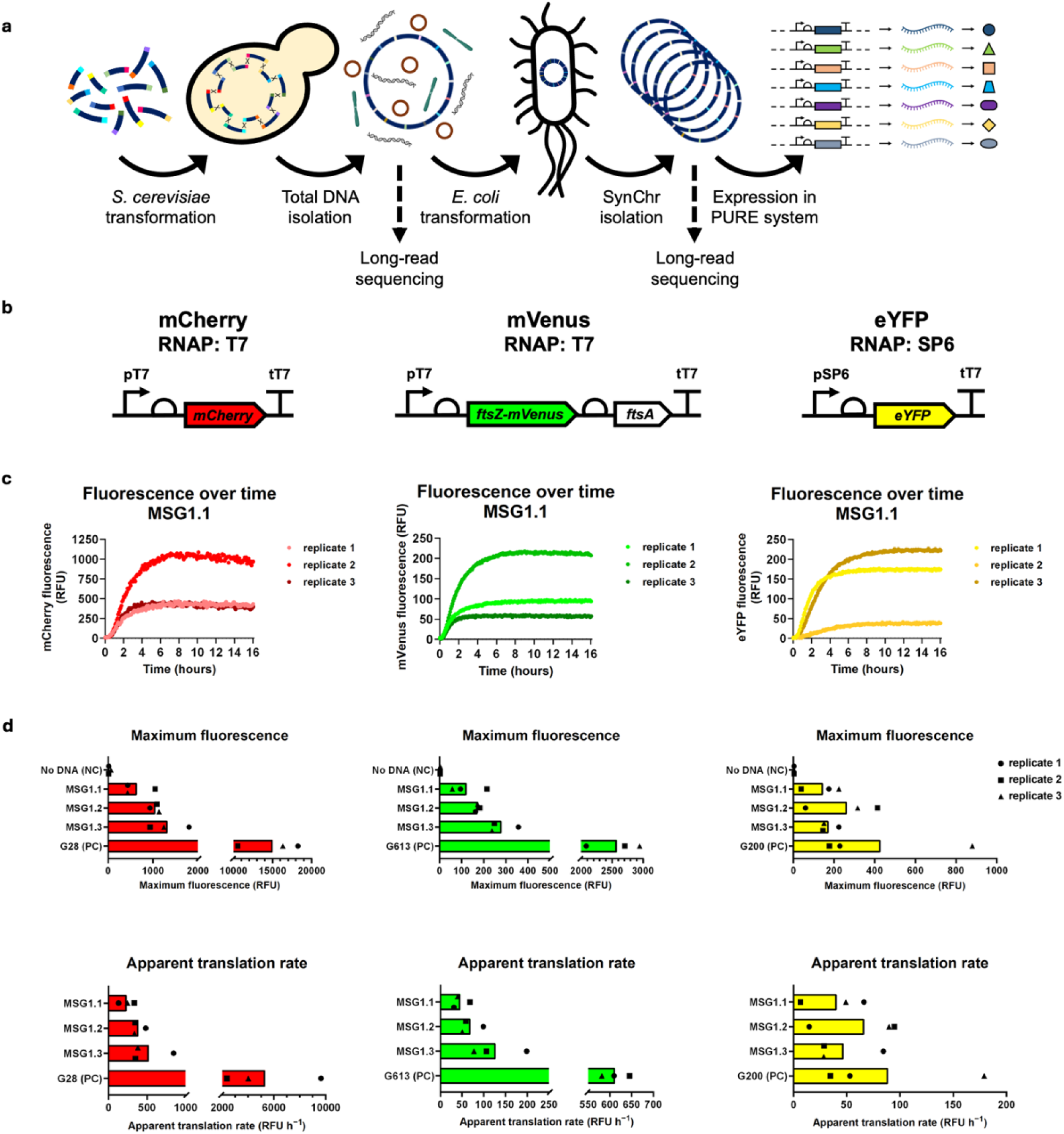
Expression kinetics in PURE system of fluorescent reporter proteins encoded on MSG1 after isolation from *E. coli*. **a)** Pipeline from assembly in yeast to expression in PURE system, involving an intermediate step for amplification of MSG1 in *E. coli*. After transformation of yeast with assembly DNA fragments, MSG1 was assembled by homologous recombination. Total DNA was extracted from yeast, verified by long-read sequencing, and *E. coli* was transformed, which selectively amplified MSG1. MSG1 DNA was isolated from *E. coli*, checked by long-read sequencing, and tested for expression in PURE system. **b)** Schematic representation of expression cassettes of *mCherry*, *mVenus* and *eYFP*. *mCherry* and *mVenus* are controlled by pT7, *eYFP* by pSP6. **c)** Fluorescence measurements over 16 h of expression of MSG1.1 in PURE system. See Supplementary Figure 10 for graphs of other MSG1 variants and control templates. **d)** Kinetic parameters of *mVenus*, *mCherry* and *eYFP* expression in PURE system: maximum fluorescence (RFU) and apparent translation rate (RFU h^−1^). All DNA templates were added at 1 nM final concentration. Bars indicate mean values across three replicates. PC, positive control; NC, negative control.

DNA concentrations measured by Qubit ranged from ca. 200 to 450 ng µL^−1^ (ca. 8–16 nM) in 50 µL. Nanodrop analysis demonstrated A_260_/_280_ ratios of 1.8–1.9 (within the desired range) and A_260_/_230_ ratios of 1.6–1.8 a (slightly below the preferred 1.8–2.2 range). Sequencing revealed contamination with *E. coli* genomic DNA at 51% for MSG1.1, 53% for MSG1.2 and 35% for MSG1.3, likely due to the low SynChr copy number ^44^. Consequently, Qubit overestimated SynChr DNA concentration by a factor two to three.

To reduce contamination by *E. coli* genomic DNA and increase the amount of isolated SynChr DNA, a high-copy number of MSG1.1 was induced by incubation with L-arabinose. This additional step yielded higher SynChr DNA concentrations (> 1000 ng µL^−1^) with reduced genomic DNA contamination (below 10%, single replicate). For this isolation, cells were grown from glycerol stock, streaked to obtain single colonies, and verified by colony PCR before being cultured for SynChr isolation. This process ensured that any potential recombination caused by cell recovery from storage was detected, allowing for SynChr isolation only from cells with non-recombined SynChrs.

### Synthesis of MSG1-encoded proteome in PURE system

To assess the expression of the three fluorescent proteins encoded by MSG1, bulk PURE reactions were performed for all three MSG1 variants. Production of mCherry and mVenus controlled by pT7, and eYFP controlled by pSP6, was tested using reactions containing either T7 RNAP or SP6 RNAP (Figure 4b). Fluorescence kinetics were measured during 16 hours. As positive controls, plasmids encoding individual fluorescent markers—originally used as templates for PCR to obtain the assembly fragments—were used. All DNA templates were added at 1 nM final concentration (for MSG1 variants, the concentrations measured by Qubit were taken, not corrected for *E. coli* genomic DNA contamination).

All fluorescent proteins were successfully expressed from the three MSG1 variants (Figure 4c and Supplementary Figure 10). Orthogonality of the SP6 RNAP and T7 promoters was confirmed (Supplementary Figure 11). Two kinetic parameters of *mVenus*, *mCherry* and *eYFP* expression were further analyzed: the apparent translation rate and maximum fluorescence. The apparent translation rate (RFU h^−1^) was estimated using sigmoidal fitting on the fluorescence data ^45^ and corresponds to the maximum slope in the linear regime of expression (Supplementary Figure 10). Maximum fluorescence (RFU) was used as a proxy for the total amount of fluorescent protein produced and was determined by identifying the 100-minute time window with the highest mean fluorescence.

Both kinetic parameter values were similar for all three MSG1 variants and lower than for the positive control templates with T7p (mCherry and mVenus). However, no strong difference was observed for eYFP, which can be explained by the fact that MSG1 contains only three expression cassettes controlled by pSP6 and eleven by pT7.

Expression of all 15 (fluorescent and non-fluorescent) proteins from the MSG1 variants was validated by mass spectrometry analysis of the same samples used for fluorescence measurements. Proteolytic peptides could be detected for all proteins under pT7 for all MSG1 variants, albeit with varying coverage levels (Figure 5a). As expected from the orthogonality between pT7 and pSP6, proteins PgsA and PgpA under control of pSP6, were not detected in these conditions. Principal component analysis of relative protein abundances from biological replicates showed clustering of all replicates of each condition and of the three MSG1 variants, indicating that relative abundance profiles were consistent and reproducible (Figure 5b). The orthogonality between T7 and SP6 RNA polymerases and their respective promoters was visualized in a volcano plot (Figure 5c). Specifically, proteins under control of pSP6 (PgsA and PgpA) were only expressed with SP6 RNAP, while proteins controlled by pT7 (all other proteins) were significantly more abundant in the presence of T7 RNAP. We then compared the raw abundances of proteins under transcriptional control of pT7 across MSG1 variants and control plasmids that only carry the genes of individual modules (Figure 5d, Supplementary Figure 12). Overall, protein abundance was higher for control plasmids than for MSG1 variants (by a factor 2 to 9), as expected from the fewer number of encoded genes.

**Figure 5:**
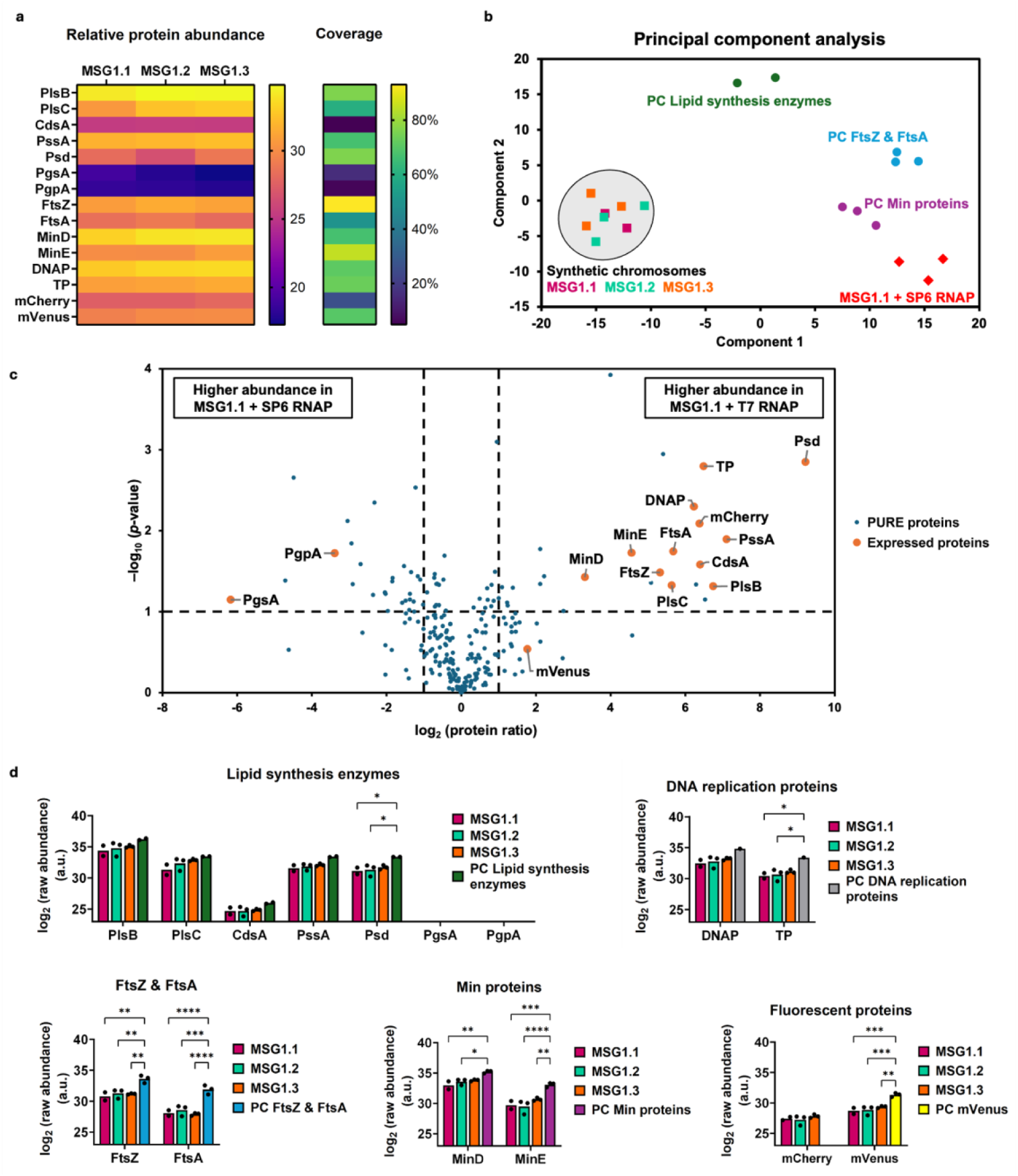
Mass spectrometry reveals protein synthesis from the MSG1 variants. **a)** Relative abundance of the MSG1-encoded proteins expressed in PURE system using T7 RNAP. The percentage of protein sequence coverage was calculated by dividing the number of amino acids in all detected peptides by the total number of amino acids in the entire protein. **b)** Principal component analysis of relative protein abundance using T7 RNAP or SP6 RNAP. **c)** Volcano plot displaying the change in relative protein abundance between MSG1.1 transcribed either by T7 RNAP or SP6 RNAP. Vertical lines indicate a 2-fold increase. The horizontal line indicates a *p*-value = 0.1 from a two-tailed *t*-test. **d)** Raw abundance of the proteins expressed with T7 RNAP, classified by functional modules. Bars indicate mean values. Individual values (n) are plotted as dots. Asterisks denote statistically significant differences per protein between DNA templates from a two-way ANOVA with Tukey’s multiple comparisons test (*, *p* ≤ 0.05; **, *p* ≤ 0.01; ***, *p* ≤ 0.001; ****, *p* ≤ 0.0001), all other differences are non-significant. *n* = 1 for the control plasmid for DNA replication proteins, *n* = 2 for MSG1.1 with T7 RNAP and the control plasmid for lipid synthesis enzymes, and *n* = 3 for all other conditions. PC, positive control.

In summary, these results show that all MSG1-encoded genes can be expressed, they confirm the orthogonality of SP6 and T7 transcriptional control, and demonstrate that changes in the type of RNAP can shift the overall proteomic output of the SynChr.

### Expression of MSG1 inside liposomes

To emulate gene expression in a synthetic cell, PURE reactions with 0.1 nM MSG1.1 were compartmentalized in liposomes and gene expression directed by T7 RNAP was monitored by fluorescence confocal microscopy (Figure 6a–c). Liposomes exhibiting fluorescence from both mVenus and mCherry proteins were observed (Figure 6b, c), demonstrating that compartmentalized expression of a 15-gene genome is feasible. Heterogeneity in expression levels across liposomes is inherent to the vesicle formation protocol and is also observed with the smaller control plasmids (Figure 6b). Fluorescence intensity of both reporter proteins in single liposomes is on average lower from MSG1 than from the control plasmids, as expected from the higher number of transcription cassettes in the SynChr.

**Figure 6:**
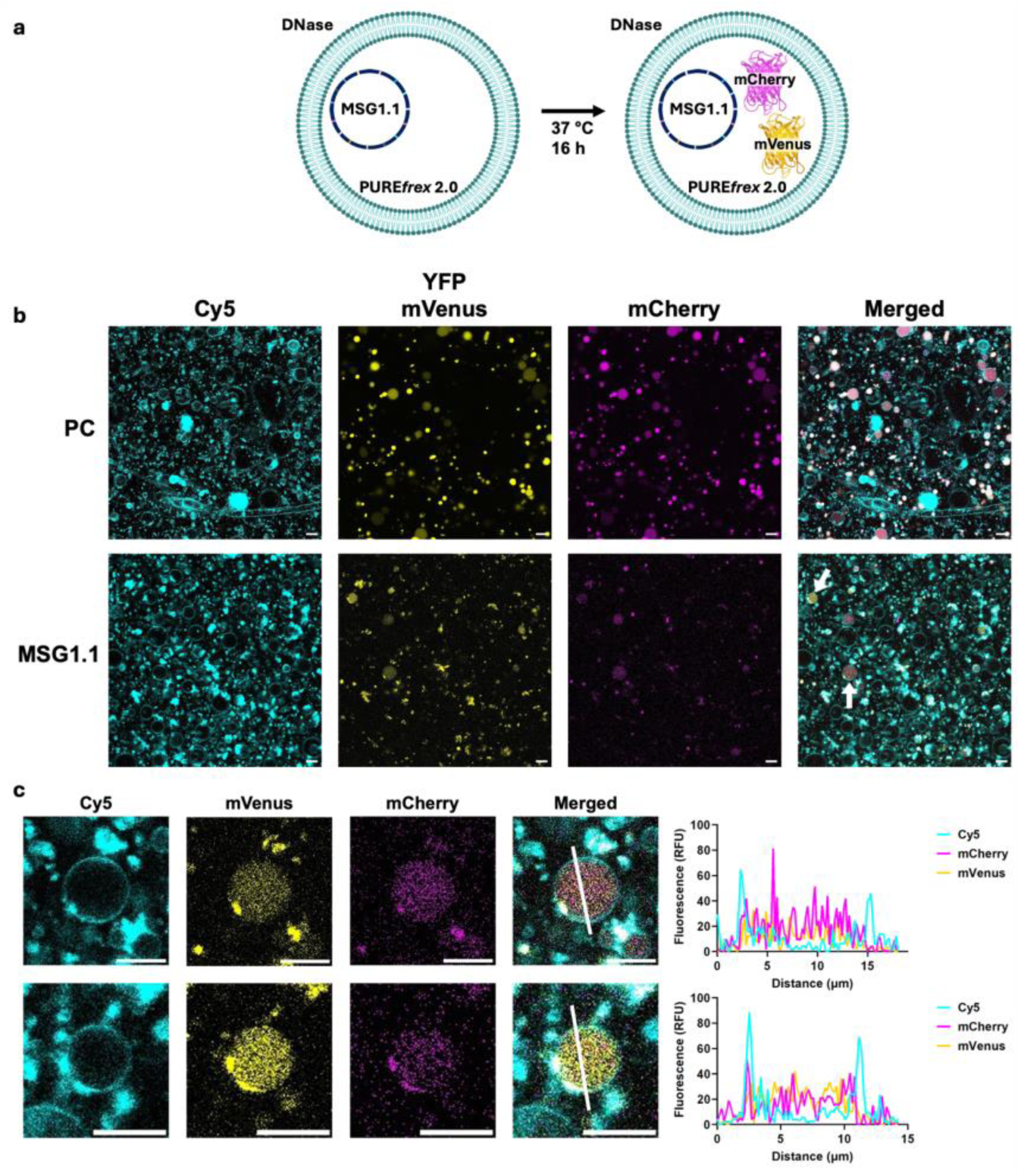
In-liposome expression of fluorescent reporter genes from MSG1.1. **a)** Schematic illustration of MSG1 expression in liposomes, directed by T7 RNAP. The circular MSG1.1 was encapsulated together with PURE system. DNase was added to prevent expression outside liposomes. Membrane dye: Cy5. Expected markers to be expressed: mVenus and mCherry. **b)** Confocal microscopy images of liposomes containing either control templates (PC, encoding *YFP* and *mCherry*) or MSG1.1. Liposomes indicated with a white arrow in the merged channel images are highlighted in c). **c)** Example liposomes showing mCherry and mVenus signals in the lumen. Fluorescence intensity profiles of Cy5, mCherry and mVenus signals were measured along the white line appended in the merged image. Scale bars are 10 µm.

### Linear MSG1 can be amplified by the φ29 DNA replication machinery

A self-replicating minimal cell must be able to replicate its genomic DNA. The φ29 system for protein-primed replication has previously been reconstituted in vitro for amplification of synthetic DNA templates up to 10 kb ^46^ and of the 19.3-kb φ29 genome itself ^38^. However, it has not been tested yet for large synthetic templates exceeding the size of the φ29 genome. The circular MSG1.1 was digested with PmeI to generate a linear template flanked by the replication origins oriL and oriR (Figure 7a). The linear MSG1 was added in PURE system (without ribosomes) supplemented with purified DNA replication proteins DNAP, TP, SSB and DSB. To distinguish newly synthesized DNA from the initial template, dTTP was partly substituted by fluorescein-dUTP and the reaction products were visualized on agarose gel. Full-length synthesis of MSG1.1 strands was confirmed by the presence of a band around 40 kb in the fluorescein channel (Figure 7c and Supplementary Figure 13). This result suggests the ability of the reconstituted φ29 system for replicating multigene DNA templates, even under transcription-translation conditions.

**Figure 7:**
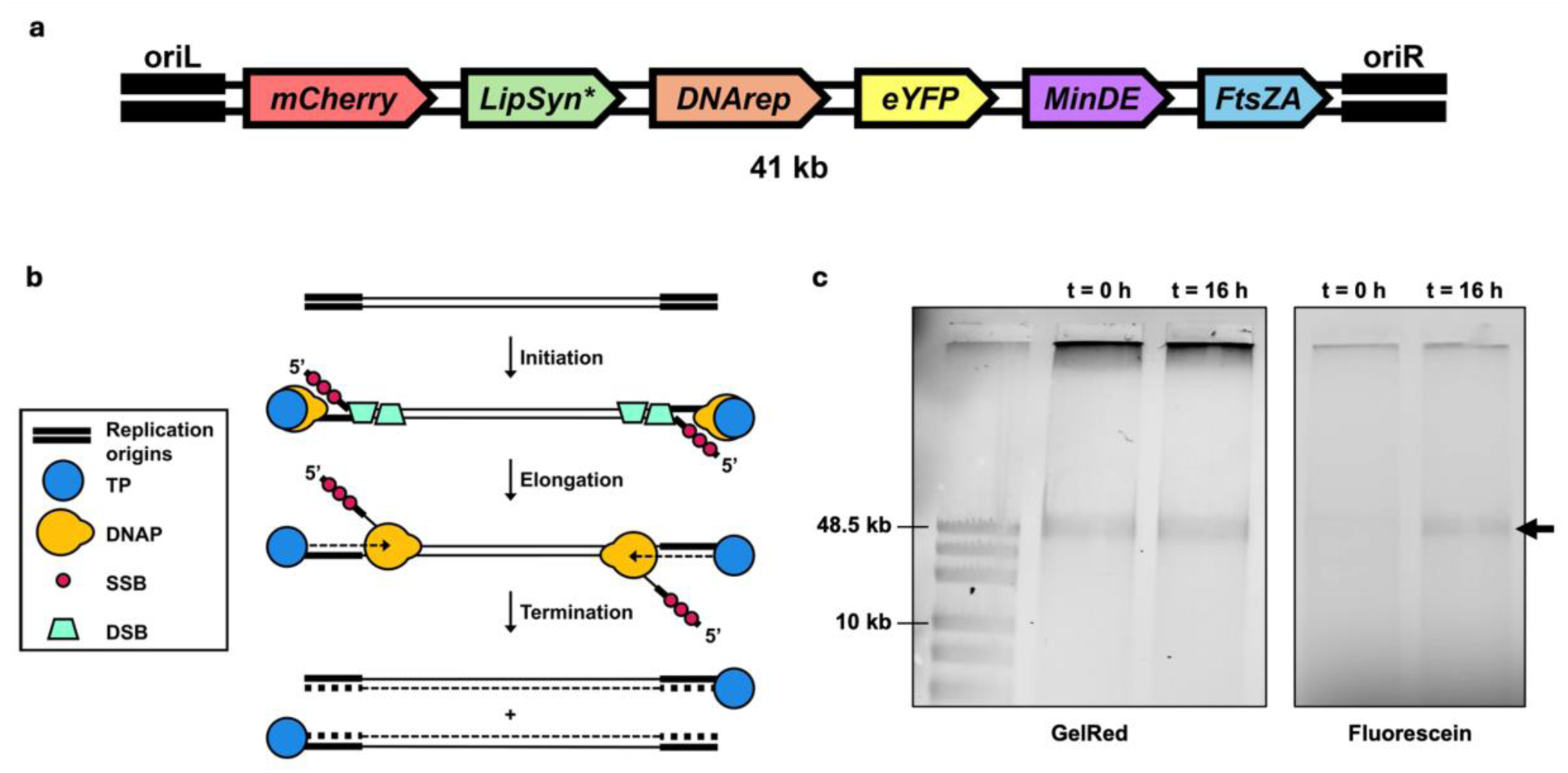
Protein-primed replication of linearized MSG1. **a)** Schematic representation of MSG1 linearized by PmeI. The order and direction of the minimal cell modules, including fluorescent reporters, is shown. oriL, left origin of replication; LipSyn, lipid synthesis enzymes; DNArep, DNA replication proteins; MinDE, MinD and MinE; FtsZA, FtsZ:mVenus and FtsA; oriR, right origin of replication. *The two lipid synthesis genes under pSP6 control (*pgsA* and *pgpA*) are encoded in the opposite direction. **b)** Schematic illustration of the protein-primed DNA replication system of bacteriophage φ29, requiring a linear template flanked with replication origins. TP, terminal protein; DNAP, DNA polymerase; SSB, single-stranded DNA binding protein; DSB, double-stranded DNA binding protein. Illustration adapted from ^38,46^. **c)** Full-length synthesis of MSG1.1 visualized on agarose gel. No ribosomes were added to the reaction. The GelRed channel shows all DNA, the fluorescein channel shows newly synthesized DNA. The arrow points at the expected band for linear MSG1.1 with a size of 41 kb.

## Discussion

A 15-cistron minimal synthetic genome (MSG1) encoding multiple (viral and bacterial) proteins involved in synthetic cell modules was successfully assembled in yeast and extracted. In vitro transcription and translation of MSG1 using PURE system enabled the one-pot synthesis of the 15 encoded proteins, demonstrating the feasibility to design, build, and express large multigene constructs encoding cellular functions for a synthetic cell. Additionally, encapsulation and expression of MSG1 in liposomes was possible, and polymerization of full-length MSG1 by the protein-primed DNA replication system of φ29 was demonstrated.

Although homologous recombination events occurred more frequently between SHRs than between the PURE repeats, undesired recombination events reduce the rate of correctly assembled SynChrs. Improving the assembly efficiency will be indispensable to scale up the number of genes and build an entire synthetic cell genome (see below). Possible solutions can be envisaged, either separately or in combination: (i) reducing the degree of similarity between PURE repeats ^47^, for instance by using existing libraries of T7 promoters ^48^ and RBS sequences ^49^, which can simultaneously serve to modulate expression levels, (ii) moving the PURE repeats further inward of the fragments to be overlooked by the HDR machinery, which predominantly targets the fragment ends, (iii) altering the processivity of the nucleases Exo1 and Sgs1-Dna2 to shorten the ssDNA overhang, excluding the PURE repeats from involvement in recombination ^50^, and (iv) using *URA3* for selection upon transformation instead of hygromycin resistance.

Current DNA isolation protocols from yeast led to suboptimal concentration and purity of centromeric SynChrs for direct expression in PURE system. The use of a 2μ replication origin to increase the copy number of SynChrs in yeast is precluded by the low assembly efficiency. We found that the pCC1BAC backbone with a tunable copy number enabled the amplification of MSG1 in *E. coli*, its maintenance, and subsequent isolation, even at low copy number (i.e., no induction). Specifically, 8–16 nM MSG1 with 35–55% *E. coli* genomic DNA contamination could be obtained at a single plasmid copy, which could be increased to over 35 nM with ca. 9% contamination when the high-copy number origin was activated. This was sufficient to conduct PURE reactions with a final DNA concentration of 1 nM, which supported expression of all encoded genes. Yet, the intermediate amplification step in *E. coli* may not be universally suitable for all SynChrs due to the unpredictable toxicity of heterologous sequences ^51,52^ or recombination ^15^.

Expression of all MSG1-encoded genes in PURE system was verified by proteomics analysis. The proteomic output could be regulated by controlling gene expression with the orthogonal T7 and SP6 RNA polymerases, which will be relevant to endow synthetic cells with regulatory mechanisms. Moreover, even if the regulatory sequences of most genes are similar, 3’ and 5’ UTR sequences vary between genes, which may lead to different protein expression levels ^53^. Stronger control over the expression of multiple genes might be achieved by implementing in PURE system the *E. coli* RNA polymerase in combination with various σ factors ^54,55^, which may also increase the assembly efficiency by decreasing the similarity of promoter sequences. Genome-wide modifications of coding or regulatory sequences may directly be implemented in yeast with CRISPR-based tools ^56^ or inside *E. coli* using recombineering methods ^57^. Furthermore, combinatorial assembly in yeast of DNA fragment library could be used to change the cistron order or orientation. All these strategies aim at creating genetic diversity, which, when combined with screening or selection of the genome variants, will establish the basis for directed evolution of system’s level functions ^11^.

MSG1 can be seen as the scaffold for the design of larger and more complete genomes for a minimal synthetic cell. Additional genes can be introduced in new assembly fragments or directly into the MSG1’s landing pad by CRISPR-mediated integration in yeast ^58^. Primary gene candidates to empower synthetic cells with a replicative core are those involved in the self-regeneration of PURE system: T7 RNA polymerase, *E. coli* translation factors ^15^, ribosome ^59^, and transfer RNAs ^60^. Finally, the ability of the φ29 machinery to replicate such a large synthetic genome (> 150 kb) remains to be explored.

## Methods

### Strains and culture conditions

*Saccharomyces cerevisiae* strains used and constructed in this study (Supplementary data 1.1 and 1.2) are derived from the CEN.PK lineage ^61^. Strains were propagated in complex medium (Yeast extract Peptone Dextrose, YPD) containing 10 g L^−1^ Bacto yeast extract (Gibco, Thermo Fisher Scientific, Waltham, MA, USA), 20 g L^−1^ Bacto peptone (Gibco) and 20 g L^−1^ glucose or in synthetic medium (Synthetic Medium Dextrose, SMD) containing 3 g L^−1^ KH_2_PO_4_, 0.5 g L^−1^ MgSO_4_·7H_2_O, 5 g L^−1^ (NH_4_)_2_SO_4_, 20 g L^−1^ glucose, trace elements and vitamins ^62^. Both media were initially prepared without glucose, set to pH 6.0 using 2 M KOH and sterilized (20 min at 110 °C for YP, 20 min at 121 °C for SM). Glucose was sterilized separately at 110 °C for 20 min before addition to the sterilized media. Filter-sterilized vitamins were added to SMD after media sterilization. For selection based on the dominant marker *hphNT*1, YPD was supplemented with hygromycin B (Hyg) to 200 mg L^−1^. For selection based on auxotrophic markers, SMD was supplemented with separately sterilized solutions of uracil (Ura) to 150 mg L^−1^, histidine (His) to 125 mg L^−1^ and/or leucine (Leu) to 500 mg L^−1^. Yeast strains were grown aerobically in liquid media at 30 °C and 200 rpm in an Innova incubator shaker (New Brunswick Scientific, Edison, NJ, USA) or on solid media plates at 30 °C.

*Escherichia coli* strains XL1-Blue (Agilent, Santa Clara, CA, USA), TOP10 (Thermo Fisher Scientific) or DH5α (Thermo Fisher Scientific) were used for molecular cloning and propagation of conventional plasmids. *E. coli* strains DH10B (Invitrogen, Thermo Fisher Scientific) and EPI300 (LGC Biosearch Technologies, Hoddesdon, UK) were used for the construction and propagation of constructs with the CopyControl pCC1BAC vector backbone ^63^. SynChrs that were assembled in *S. cerevisiae* contained a pCC1BAC vector backbone and were propagated in EPI300 after isolation from yeast. *E. coli* strains were grown in Lysogeny Broth (LB) medium containing 5 g L^−1^ Bacto yeast extract, 10 g L^−1^ Bacto tryptone (Gibco) and 5 g L^−1^ NaCl, supplemented with 50–100 mg L^−1^ ampicillin (Amp), 50 mg L^−1^ kanamycin (Kan) or chloramphenicol (Cam, 25 mg L^−1^ for high-copy number plasmids, 12.5 mg L^−1^ for constructs with pCC1BAC backbone) if required. *E. coli* strains were grown at 37 °C in 10 mL liquid medium in 50 mL centrifuge tubes at 250 rpm in an Innova 4000 incubator shaker (New Brunswick Scientific), in 100 mL liquid medium in 500 mL shake flasks at 200 rpm in an Innova 44 incubator shaker or on solid media plates at 37 °C.

Solid media were prepared by adding 20 g L^−1^ Bacto agar (Gibco) to the medium prior to heat sterilization. All *S. cerevisiae* and *E. coli* strains were stored at −80 °C in aliquots of overnight grown culture supplemented with sterile 30% (v/v) glycerol.

### PCR and clean-up

Amplification of *Arabidopsis thaliana* genomic DNA (gDNA) fragments was performed using the LongRange PCR Kit (Qiagen, Venlo, The Netherlands) according to the manufacturer’s instructions.

Amplification of DNA fragments for plasmid construction was performed using KOD Xtreme Hot Start DNA Polymerase (Sigma-Aldrich, Merck, Darmstadt, Germany) according to the manufacturer’s protocols, or using Phusion High-fidelity DNA Polymerase (Thermo Fisher Scientific) according to the manufacturer’s instructions or with a reduced primer concentration of 0.2 µM. Fragments for assembly in yeast (Supplementary data 4.2 and 4.3) were obtained with Phusion, KOD Xtreme or UltraRun LongRange PCR kit (Qiagen, Venlo, The Netherlands), following the manufacturer’s instructions except that we lowered the primer concentration to 0.2 µM for Phusion reactions and scaled up the reaction volume to 50 µL for UltraRun LongRange reactions. In case of unsuccessful amplification, primer-dimer formation was reduced by lowering the primer concentration tenfold, addition of 5% DMSO, performing an initial denaturation step for 3 min and one in the cycling stage for 20 s. Verification of constructed plasmids in *E. coli* was done through colony PCR using GoTaq DNA polymerase (Promega, Madison, WI, USA) according to the manufacturer’s instructions, directly using a resuspended colony or with the addition of an extra lysis step. In the latter case, a colony was resuspended in 50 µL Milli-Q and cells were lysed at 95 °C for 5 min, after which 2.5 µL of lysed cells was used in the PCR reaction. Alternatively, plasmids and SynChrs in *E. coli* were verified by colony PCR using DreamTaq PCR Master Mix (Thermo Fisher Scientific) according to the manufacturer’s instructions, downscaled to 10 µL reaction volume, using a resuspended colony and with an initial incubation at 95 °C for 10 min to lyse the cells and release the DNA.

Primers used to generate fragments for assembly in yeast were PAGE-purified and purchased from Sigma-Aldrich. All other primers were desalted when shorter than 30 nt and HPLC-purified or PAGE-purified when longer. IMB primers were purchased from Sigma-Aldrich, ChD primers from Ella Biotech (Fürstenfeldbruck, Germany) or Biolegio (Nijmegen, The Netherlands). All primers are listed in Supplementary data 3.

PCR product size was analyzed by agarose gel electrophoresis on a 0.6–1 % agarose gel in 1× TAE buffer, with the GeneRuler DNA Ladder Mix (Thermo Fisher Scientific), BenchTop 1kb DNA Ladder (Promega) or Quick-Load 1 kb Extend ladder (New England Biolabs) as reference. Fragments used for plasmid construction were treated with DpnI (Thermo Fisher Scientific or New England Biolabs) to remove the parental plasmid, by addition of 1 µL DpnI enzyme directly to the 50 µL PCR sample and incubation at 37 °C for 30 min. Purification of PCR products used for cloning was done using InnuPREP PCRpure Kit (IST Innuscreen GmbH, Berlin, Germany) according to the manufacturer’s instructions with additional centrifugation for 2 min after discarding the binding buffer, using the Wizard SV Gel Clean-Up System (Promega) following the manufacturer’s protocol or using the GeneJET PCR purification kit (Thermo Fisher Scientific) according to the manufacturer’s instructions. In case of undesired PCR side products, purification from agarose gel was performed using ZymoClean Gel DNA Recovery kit (Zymo Research, Irvine, CA, USA) or the Wizard SV Gel and PCR Clean-Up System (Promega) according to the manufacturer’s instructions. For all kits, DNA was eluted after 5–10 min incubation with 50–60 °C Milli-Q.

Fragments for assembly in yeast of MSG0.1 and MSG0.2 (Supplementary data 4.2) were obtained by pooling multiple PCR reactions, treatment with DpnI and purification using the GeneJet PCR purification kit.

All fragments for assembly in yeast of MSG1 (Supplementary data 4.3) were obtained by pooling eight to eleven 50 µL PCR reactions, treatment with DpnI and purification from gel to minimize the chance of transforming yeast with parental plasmids or side products. Extraction from gel was done using the ZymoClean Gel DNA Recovery kit according to manufacturer’s instructions with the following modifications: the columns were incubated with the melted agarose solution for at least 30 min before centrifugation, three washes were performed with 2 min incubation of the wash buffer, residual ethanol was removed by 2 min centrifugation of the column after removal of the flowthrough in the last wash step, and the column was incubated for 5–10 min with 50–60 °C Milli-Q prior to elution. Alternatively, ZymoClean Large Fragment DNA Recovery Kit (Zymo Research) was used according to the manufacturer’s instructions with the modifications described above, or the QIAquick Gel Extraction Kit (Qiagen) was used according to manufacturer’s protocol with an additional wash step with Buffer PE, incubation of the column with Buffer PE for 2 min prior to centrifugation, and 5 min incubation with 50–60 °C Milli-Q before elution of the DNA.

### In vitro plasmid assembly

Small insertions or mutations were introduced in plasmids using site-directed mutagenesis PCR, where the mutation or insertion was included in the overhang of the primer. The PCR product was circularized in one of the following two ways: (i) Phosphorylated primers were used during PCR and after DpnI digestion and optionally cleanup, the PCR product was self-ligated by T4 ligase (Thermo Fisher Scientific), following the manufacturer’s instructions. (ii) Primers without phosphorylation were used during PCR and after DNA extraction from gel, the fragment was circularized by Gibson assembly, following the third protocol described below.

Gibson assembly of one or multiple linear fragments was performed in one of the following three ways. (i) Using the NEBuilder HiFi Master Mix (New England Biolabs) according to the manufacturer’s protocol, downscaled to 5 µL or 10 µL reaction volume and with an incubation of 1 h at 50 °C. (ii) Using the Gibson Assembly Master Mix (New England Biolabs) according to the manufacturer’s protocol. (iii) Using a protocol adapted from ^64^: A 5× ISO buffer (0.5M Tris-HCl pH 7.5, 50 mM MgCl_2_, 1 mM of each of the four dNTPs, 50 mM DTT, 25% PEG-8000, 5 mM NAD) was prepared according to the protocol in ^64^. An adapted Master Mix was assembled, containing 320 µL 5× ISO buffer, 0.64 µL 10 U µL^−1^ T5 exonuclease, 20 µl 2 U µL^−1^ Phusion Polymerase, 160 µL 40 U µL^−1^ Taq ligase and 700 µL Milli-Q. One 20 µL Gibson Assembly reaction consisted of 15 µL Master Mix with 100 ng of the linear PCR product in Milli-Q. The reaction was incubated at 50 °C for 60 min.

Golden Gate assembly of the template plasmids for MSG0.1 and MSG0.2 was performed according to the protocol described in ^65^ with adjustments. For level 0 assemblies, insert concentration was increased to 40–150 fmol at an entry vector concentration of 20 fmol, and 30 cycles of digestion and ligation were performed. For level 1 assemblies, digestion in each cycle was carried out at 37 °C for 3 min. For all other Golden Gate reactions, the reaction volume was downscaled to 5 µL and all digestion steps were performed at 37 °C.

### *E. coli* transformation with plasmids and SynChrs

*E. coli* was transformed with in vitro assembled plasmids for propagation and storage, following one of the protocols described below.

*E. coli* XL1-Blue cells (Agilent), made chemically competent in-house according to the protocol described in ^66^, were used for transformation of ligated site-directed mutagenesis PCR products and of Golden Gate reactions. A mixture of 5 µL ligation product or Golden Gate reaction and 50 µL cells was incubated on ice for 5 min. Cells were transformed via heat shock for 45 s at 42 °C, followed by incubation on ice for 2 min and resuspension in 450 µL Super Optimal broth with Catabolic repression (SOC), containing 5 g L^−1^ Bacto yeast extract, 20 g L^−1^ Bacto tryptone, 0.58 g L^−1^ NaCl, 0.19 g L^−1^ KCl, 2 g L^−1^ MgCl_2_·6H_2_O, 2.46 g L^−1^ MgSO_4_·7H_2_O and 3.6 g L^−1^ glucose. The cells were incubated for 1 h at 37 °C and 250 rpm, plated on LB agar plates with appropriate antibiotics and grown overnight at 37 °C.

Chemically competent *E. coli* TOP10 cells (Thermo Fisher Scientific) were used for transformation of site-directed mutagenesis PCR products that were circularized by Gibson assembly. To 50 µL of cells, 5 µL of Gibson assembly mix was added and the cells were incubated for 10 min on ice. Heat shock was performed as described above, but replacing 450 µL SOC by 1 mL LB medium. For G131 cloning, 1 µL of PCR product was mixed with 50 µL TOP10 chemically competent *E. coli* and transformation was performed as described above for XL1-Blue cells.

Chemically competent *E. coli* DH5α cells (Thermo Fisher Scientific) were used for transformation of Gibson Assembly products obtained using the Gibson Assembly Master Mix. The transformation protocol was the same as for TOP10 cells.

*E. coli* DH10B cells (Invitrogen) were used for electroporation of NEBuilder Hifi assembly products containing a pCC1BAC backbone. The cells were made electrocompetent in-house by growing them in 300 mL LB without NaCl until an OD_600_ of 1.6, incubation on ice for 10 min, and performing multiple rounds of centrifugation for 10 min at 4000 g and 4 °C followed by resuspension of the cell pellet in 150 mL, 40 mL, 15 mL and 600 µL ice-cold 10% glycerol, respectively. The electrocompetent cells were stored in 50 µL aliquots at −80 °C until use. For transformation, 2 µL of NEBuilder Hifi assembly product was mixed with 50 µL electrocompetent cells and electroporation was performed with the MicroPulser Electroporator (Bio-Rad Laboratories, Hercules, CA, USA) according to the manufacturer’s instructions, using 0.2 cm gap Gene Pulser/MicroPulser Electroporation cuvettes (Bio-Rad Laboratories).

*E. coli* TransforMax EPI300 cells (LGC Biosearch technologies) were used as host for the propagation and storage of SynChrs assembled in yeast, which contain a pCC1BAC backbone. Following extraction of total cellular DNA from yeast, 1 µL DNA was mixed with 50 µL cells and electroporation was performed as described for DH10B. Since the SynChr is the only DNA in the total cellular DNA of yeast that contains a pCC1BAC backbone, it was expected that transformation of *E. coli* results in selective amplification of SynChrs.

### Plasmid and SynChr isolation from *E. coli* and verification

Plasmid isolation from *E. coli* was performed using the PureYield Plasmid Miniprep System (Promega) or the GeneJET Plasmid Miniprep Kit (Thermo Fisher Scientific), according to the manufacturer’s instructions. Elution was done in 25 µL 50–60 °C Milli-Q.

Plasmids isolated from *E. coli* were verified by one or multiple of the following methods: (i) colony PCR as described above, (ii) restriction analysis using FastDigest enzymes (Thermo Fisher Scientific) or CutSmart enzymes (New England Biolabs), according to the manufacturer’s protocol, and visualization with agarose gel electrophoresis, (iii) Sanger sequencing by Macrogen Europe (Amsterdam, The Netherlands) or (iv) whole-plasmid sequencing using Oxford Nanopore technology by Plasmidsaurus (Eugene, OR, USA).

SynChr isolation from EPI300 *E. coli* cells was performed using the NucleoBond Xtra Midi kit (Macherey-Nagel, Düren, Germany). No induction of the high-copy number origin on the pCC1BAC backbone was performed, so the SynChr was expected to be maintained at one copy per cell. After cultures were grown directly from colonies on the transformation plate, the resulting isolated DNA consisted of a mixture of recombined and non-recombined SynChrs, as determined by whole-plasmid sequencing. Therefore, extra care was taken to ensure that a single species of SynChr was isolated. Colonies from the transformation plate were streaked to obtain single colonies. Single colonies were resuspended in 15 µL Milli-Q and 1 µL was used per colony PCR reaction to check for the presence of three marker fragments containing the pCC1BAC backbone, *mRuby2* gene and CEN6/ARS4 sequence, using primer pairs 15812/20422, 20333/2306 and 20641/3232, respectively (Supplementary data 3.4). The same resuspended colony was used to inoculate 10 mL LB with 12.5 µg mL^−1^ chloramphenicol and the culture was incubated overnight at 37 °C without shaking. The next morning, the 10 mL culture was used to inoculate 300 mL LB with 12.5 µg mL^−1^ chloramphenicol, which was incubated overnight at 37 °C and 200 rpm. Of the final culture, 1–2 mL was used to prepare a glycerol stock. The remaining culture volume was utilized for SynChr isolation with the NucleoBond Xtra Midi kit, starting from an OD_600_ × culture volume (mL) of 800. The manufacturer’s instructions were used for purification of low-copy plasmids with the following modifications: extra care was taken to mix gently after addition of Buffer NEU to prevent formation of clouds and thereby clogging the filter-column combination, 15 µL GlycoBlue (Invitrogen) was added after elution with Buffer Elu to ease visualization of the DNA pellet, centrifugation after addition of isopropanol was extended to 45 min, and the DNA pellet was dissolved by addition of 50 µL 50–60 °C Milli-Q and incubation overnight at 4 °C.

Another isolation was performed with high-copy number induction. Single colonies were obtained from the glycerol stock by streaking directly on an agar plate. Colonies were checked by colony PCR and grown overnight in LB with chloramphenicol as described above. The next morning, 9 mL of overnight culture was used to inoculate 291 mL LB with 12.5 µg mL^−1^ chloramphenicol and 300 µL of CopyControl solution (LGC Biosearch Technologies) was added. After overnight incubation at 37 °C and 200 rpm, SynChr isolation was performed as described above.

SynChrs sequenced by Plasmidsaurus as big plasmid were analyzed by looking at (i) the consensus sequence, (ii) histogram with read lengths, and (iii) raw reads. The consensus sequence was aligned in SnapGene (version 7.0.2, GSL Biotech, San Diego, CA, USA) to the designed SynChr sequence to check for deletions and point mutations. The histogram was analyzed to determine whether a mixture or a single configuration of SynChrs was present. For analysis of the raw reads, a reference FASTA file containing the sequence of CENPK113-7D ^67^ concatenated with the designed SynChr sequence was prepared. An annotation (.gff) file of the designed SynChr was made by exporting the annotations from SnapGene and using the R package labtools (version 0.1.0, function: *make_gff_from_snap*) ^68^ in R Statistical Software (version 4.1.2) ^69^. The raw reads were mapped with minimap2 (parameters: *-ax map-ont*) ^70^ to the reference FASTA file and were sorted and indexed using SAMTools ^71^. Visualization of the mapped reads was done in the Integrated Genomics Viewer (IGV, version 2.11.9) ^72^.

Raw reads and consensus sequences are provided in Supplementary Data 7. An overview of mutations in the MSG1 variants is provided in Supplementary data 4.5.

### *S. cerevisiae* transformation

*S. cerevisiae* transformation was performed using the high-efficiency lithium acetate/single-stranded carrier DNA/polyethylene glycol method ^73^ with the following modifications. After washing the cells with 25 mL of sterile water, cells were resuspended in 1 mL 0.1 M lithium acetate (LiAc), the cells were pelleted, the supernatant was removed and 0.1 M LiAc was added to a total volume of 500 µL. The cells were resuspended and divided in 50 µL aliquots per transformation, pelleted again, the supernatant was removed and the transformation mix was added on top of the cell pellet. The transformation mix contained 240 µL 50 % PEG 3350, 36 µL 1M LiAc, 25 µL 2 mg mL^−1^ single-stranded carrier DNA and 50 µL DNA in water (resulting in a total transformation mix volume of 351 µL), extra 30 min incubation at 30 °C was performed before heat shock for 30 min at 42 °C and the cells were incubated 1–2 h at 30 °C in 1 mL YPD before plating on YPD + Hyg agar plates.

### Total DNA extraction from *S. cerevisiae*

Total DNA extraction from *S. cerevisiae* for whole-genome sequencing and transformation of *E. coli* for SynChr propagation was performed using the Qiagen Blood & Cell Culture Kit with 100/G Genomic-tips (Qiagen) according to the manufacturer’s protocol for yeast samples.

### DNA analysis

The NanoDrop 2000 UV-Vis spectrophotometer (Thermo Fisher Scientific) was used for determination of DNA purity and concentration. DNA concentration was additionally measured with a Qubit 2.0 or 4.0 Fluorometer (Invitrogen, Thermo Fisher Scientific) using the Qubit dsDNA Broad Range Assay kit (Invitrogen). Samples containing SynChrs isolated from yeast were additionally analyzed by quantitative PCR (qPCR) on the QuantStudio 5 Real-Time PCR system (Applied Biosystems, Thermo Fisher Scientific) using the PowerUp SYBR Green Master Mix (Applied Biosystems), according to the supplier’s instructions. Data were analyzed with QuantStudio Design & Analysis software v1.5.1 (Applied Biosystems). Plasmid pUDC191 purified from *E. coli* and quantified using Qubit was used to prepare a standard curve of known concentrations ranging from 100 fM to 1 nM (Supplementary data 2.4). Primers used for qPCR are listed in Supplementary data 3.8.

### Plasmid construction in *E. coli*

All plasmids used and constructed in this study are listed in Supplementary data 2. Primers used for plasmid construction are listed in Supplementary data 3.1–3.3. Primers used for verification by *E. coli* colony PCR and Sanger sequencing are listed in Supplementary data 3.4 and 3.5, respectively.

### PURE cassette plasmids for MSG0.1 and MSG0.2 construction

Plasmids consisting of non-coding DNA fragments originating from *A. thaliana* flanked by PURE repeats (Supplementary data 2.1) were assembled via Golden Gate according to the yeast toolkit principle ^67^. Part plasmids (level 0) were constructed by PCR amplification of the target region with primers containing part type-specific overhangs and assembled into the entry vector pYTK001 (*gfp* dropout) by Golden Gate cloning. Twelve-part plasmids were assembled: a T7 promoter-lacO-g10L RBS-T7 tag sequence (*pT7*, amplified from G149) as a type 2 part plasmid, a *yfp* gene (amplified from G365) and nine 5-kb *A. thaliana* chunks (C2–C10) as type 3 part plasmids, and a T7 terminator (*tT7*, amplified from G131) as a type 4 part plasmid. All primers used for PCR amplification to create PURE cassette plasmids are listed in Supplementary data 3.1. To construct the *A. thaliana* chunk part plasmids, gDNA from *A. thaliana* ecotype Columbia (Col-0) was donated by Emma Barahona and Alvaro Eseverri (Center for Plant Biotechnology and Genomics, Madrid, Spain). Nine 5-kb fragments (“chunks”) that did not contain any BsmBI and BsaI sites were selected from the *A. thaliana* Col-0 reference genome (TAIR10) in silico. DNA chunks amplified from *A. thaliana* gDNA were obtained using the LongRange PCR kit in two sequential PCR reactions to ensure sufficient DNA yield. 2 µL of the product of the first PCR reaction with external primers was used as template for the second PCR with internal primers, which resulted in 5-kb fragments flanked with type 3 overhangs for Golden Gate cloning. For chunk 5, only the PCR with internal primers was required. Golden Gate part plasmid assemblies were transformed into *E. coli* XL1-Blue and transformants were plated on LB + Cam agar plates. For each assembly, two to eight white colonies (indicating absence of the *gfp* gene) were verified by colony PCR using primer pair 14036/19265. Level 0 plasmids containing C10 and *tT7* (pGGKp363 and pGGKp348, respectively, Supplementary data 2.1) were verified by long-read whole-plasmid sequencing using Nanopore technology by Plasmidsaurus (Eugene, OR, USA).

For the assembly of *pT7*-chunk-*tT7* cassette plasmids (level 1), six level 0-part plasmids were combined in a BsaI Golden Gate assembly reaction: (i) pYTK002 (LS connector), (ii) pGGKp346 (*pT7*), (iii) pGGKp347 (*yfp*), pGGKp350 (chunk C5), pGGKp357 (chunk C2), pGGKp358 (chunk C3) or pGGKp361-63 (chunk C7, C9 and C10), (iv) pGGKp348 (*tT7*), (v) pYTK067 (R1 connector), and (vi) pYTK095 (*gfp* dropout). The resulting assemblies were transformed into *E. coli* XL1-Blue and plated on LB + Amp agar plates. Correct assemblies were verified by colony PCR of two to eight white colonies using primer pair 14776/19088. The start and end of the cassettes were confirmed by Sanger sequencing using primers 10320 and 10325 to ensure that all cassettes contained the same PURE repeats.

Due to difficulties in the level 1 Golden Gate cloning of chunks C4, C6 and C8, cassette plasmids with these chunks were constructed via Gibson assembly. pUD1251 excluding the *yfp* gene was PCR amplified using primer pair 19525/19526. The chunks were amplified from their corresponding level 0 plasmids with primers containing 20-bp homology flanks to the pUD1251 backbone (Supplementary data 3.1). After DpnI digestion and purification using the GeneJET PCR Purification kit, Gibson assembly was carried out as described earlier. Plasmids were transformed into *E. coli* XL1-Blue and plated on LB + Amp agar plates. Assemblies were verified via diagnostic colony PCR using primer pair 14776/19088 and via Sanger sequencing with primers 10320 and 10325.

Level 1 plasmids containing C2–C10 (Supplementary data 2.1) were verified by long-read whole-plasmid sequencing using Nanopore technology by Plasmidsaurus.

### crt expression cassette plasmids

Plasmids containing the β-carotene biosynthesis genes under control of a *S. cerevisiae* promoter and terminator, namely pUD1248 (*pPGK1-crtYB-tPGK*), pUD1249 (*pHHF2-crtE-tADH1)* and pUD1250 (*pTDH3-crtI-tTDH1*), were constructed via Gibson assembly (Supplementary data 2.2). Individual parts were amplified using Phusion PCR with primers containing 20-bp overlapping overhangs for assembly. Template plasmids and primers for PCR amplification are listed in Supplementary data 2.2 and Supplementary data 3.2, respectively. Plasmid pUDE269 was used as template for amplification of *crtYB*, *crtI*, and *crtE*. As pUDE269 contains these genes in a polycistronic cassette, PCR primers were designed to add the necessary start and stop codons: a stop codon for *crtYB* and *crtI* and a start codon for *crtI* and *crtE*. The PCR products were DpnI-digested, purified and assembled in a Gibson assembly reaction. Assemblies were transformed into *E. coli* XL1-Blue and plated onto LB + Cam agar plates. Assembled plasmids were verified via restriction analysis using DraI and Sanger sequencing.

Plasmid pUD1386 (Supplementary data 2.2) was constructed through site-directed mutagenesis PCR with phosphorylated primers, self-ligation with T4 DNA ligase, and heat-shock transformation of *E. coli* XL1-Blue. It was constructed by insertion of the ARS417 sequence downstream of the *crtI* expression cassette in pUD1250 using primers 20425/20426. Verification was done by colony PCR with primer pairs 20427/20428, 20427/20430 and 20429/20428, and by whole-plasmid sequencing.

### Other plasmids

Primers used for construction of other plasmids are listed in Supplementary data 3.3.

Plasmid G131 (Supplementary data 2.1) was constructed by site-directed mutagenesis PCR on plasmid G146 to remove the DraI restriction site, using primer pair 640/641 and subsequent circularization through homologous recombination in *E. coli* TOP10. The plasmid was verified by restriction analysis using EcoRI, XbaI and DraI and by Sanger sequencing with primers 106, 107, 337, 338, 339 and 340.

Plasmids pUD1226, pUDE1110, pUDE1039 and pUDE1217 (Supplementary data 2.2) were constructed via Golden Gate assembly and transformation into *E. coli* XL1-Blue. pUD1226 was constructed with plasmids pYTK013 (*pTEF1*), pGGKp304 (*ymNeongreen*) and pYTK054 (*tPGK1*) in the pGGKd015 backbone. pUDE1110 was constructed with plasmids pYTK009 (*pTDH3*), pGGKp303 (*ymTurquoise2*) and pYTK053 (*tADH1*) in the pGGKd034 backbone. Plasmids pUD1226 and pUDE1110 were checked by colony PCR using primers 10320 & 10325. pUDE1039 was constructed with plasmids pYTK018 (*pALD6*), pGGKp302 (*ymScarletI*) and pYTK051 (*tENO1*) in the pGGKd017 backbone and verified by colony PCR using primer pairs 10320/10335 and 10320/2442. pUDE1217 was constructed from four PCR products containing BsmBI restriction sites, amplified from pUD1226, pUDE1110, pUDE1039 and pGGKd71 with primer pairs 19059/19060, 19061/19062, 19063/19064 and 19065/19066, respectively. pUDE1217 was verified by restriction analysis using NcoI and SspI and Sanger sequencing with primers 865, 1779 and 10901.

Plasmids pUDC436 and pUD1394 (Supplementary data 2.2) were constructed through site-directed mutagenesis PCR similarly to plasmid pUD1386 described above. Plasmid pUDC436 was obtained through insertion of the ARS1 sequence downstream of the *mTurquoise2* expression cassette in plasmid pUDC192 with primer pair 20431/20432. The plasmid was verified by colony PCR using primer pairs 20433/20434, 20433/20436 and 20435/20434, and its sequence was confirmed by whole-plasmid sequencing. Plasmid pUD1394 was designed to include an I-SceI cut site and a landing pad upstream of the CEN6/ARS4 sequence of pYTK081. The I-SceI cut site was included to allow linearization of the SynChrs and subsequent visualization on a CHEF gel. The landing pad was introduced to allow easy CRISPR/Cas9 editing of the SynChrs after assembly in yeast, and consists of two gRNA target sites (“Cas9 Target Site 9 (T9)” from ^41^ and “sTarget#2” from ^40^). The first site is flanked by two 60-bp non-coding sequences without homology to the yeast genome. The second gRNA target site is located downstream of the right flank, and is directly followed by the CEN6/ARS4 sequence, to allow exchange of the centromeric region. pUD1394 was constructed using primer set 20522/20523 for site-directed mutagenesis PCR on pYTK081. The insertion was verified by colony PCR with primer set 19945/20428 and the plasmid sequence was confirmed with whole-plasmid sequencing.

Plasmids G162, G197 and G200 (Supplementary data 2.3) were constructed through site-directed mutagenesis PCR, Gibson assembly to circularize the linear product following the protocol adapted from ^64^, and heat-shock transformation of *E. coli* TOP10. Plasmid G162 was constructed by replacing the DraI cut site in pETORPHI by a PmeI cut site using primer set 683/684. The modification was confirmed by Sanger sequencing using primer 288 and the plasmid sequence was determined by whole-plasmid sequencing. Plasmid G197 was obtained by insertion of a *lacO* site upstream of the *yfp* gene of plasmid G76 using primer pair 715/716 and verified by Sanger sequencing with primer 365. Plasmid G200 was constructed through replacement of *pT7* in G197 by *pSP6*, using primer set 719/720. Verification of the promoter sequence was done by Sanger sequencing with primer 365 and the plasmid was sequenced using whole-plasmid sequencing.

Plasmid pUD1387 (Supplementary data 2.2) was constructed through Gibson assembly using NEBuilder HiFi Master Mix and electroporation of *E. coli* DH10B. The *pURA3-URA3-tURA3* fragment was amplified from pYTK074 using primers 20419/20420 and was inserted into the pCC1BAC backbone, amplified from pCC1BAC-lacZα using primers 20417/20418. The constructed plasmid was verified by colony PCR with primer sets 20421/20422, 20421/20424 and 20423/20422, and by whole-plasmid sequencing.

Plasmids G607 and G613 (Supplementary data 2.3) were constructed through Gibson assembly using the Gibson Assembly Master Mix and heat-shock transformation of *E. coli* DH5α. Plasmid G607 was designed to encode a fusion protein of FtsZ with the fluorescent reporter mVenus. For construction of G607, plasmid G379 containing *pT7-ftsZ-tT7* was linearized by PCR using primers 1491/1492, thereby splitting the *ftsZ* coding sequence between glycine 55 and glutamine 56 and inserting the linkers GSTLE and LEGST downstream and upstream of each respective residue ^74^. The *mVenus* coding sequence was amplified from pWKD014 using primers 1493/1494, thereby flanking the fragment with GSTLE and LEGST linkers on its 5’ and 3’ ends, respectively. After Gibson assembly of the two fragments, the resulting G607 plasmid was verified by colony PCR using primer set 770/797. Plasmid G613 was designed to contain an operon encoding the fusion protein FtsZ-mVenus and FtsA. The *ftsA* fragment was amplified from G385 with primers 709/1508 and the resulting PCR product included a 15-bp spacer sequence with 30% GC content, generated via the Random DNA generator provided by the Maduro Lab (https://faculty.ucr.edu/∼mmaduro/random.htm), upstream of the T7 gene 10 leader sequence and RBS of the expression cassette. Plasmid G607, containing *pT7-ftsZ:mVenus-tT7*, was linearized by PCR using primers 452/1511 and contained the same 15-bp spacer sequence downstream of the *ftsZ:mVenus* stop codon. After Gibson assembly of the two fragments, correct insertion was verified by colony PCR with primer set 984/1211. The sequence of G613 was verified by whole-plasmid sequencing.

### Assembly of SynChrs in yeast

*S. cerevisiae* strain CEN.PK102-12A was used for the assembly of MSG0.1 and MSG0.2, each from 20 assembly fragments. Strain CEN.PK113-5D was used for assembly of MSG1 from 14 fragments. Host strains are listed in Supplementary data 1.1 and assembly fragments in Supplementary data 4.2 and 4.3.

Fragments for assembly of MSG0.1 and MSG0.2 include a *yfp* cassette for expression in PURE system (PF1), nine 5-kb *A. thaliana* chunks with or without PURE repeats (PF2–10 and CF2–10), a yeast replication origin (CEN6/ARS4 or 2µ) combined with an *E. coli* replication origin (ColE1) and antibiotic marker (*bla*), and nine yeast markers (*mRuby2*, *mTurquoise2*, *HIS3*, *URA3*, *LEU2*, *crtE*, *crtI*, *crtYB* and *hphNT1*). Chunk 2 harbored an I-SceI restriction site, allowing the linearization for screening purposes.

Fragments for MSG1 assembly include six fragments containing genes for expression in PURE system: the DNA replication genes *p2* and *p3*, the Min system genes *minD* and *minE*, the division protein genes *ftsZ:mVenus* and *ftsA*, the phospholipid synthesis genes *plsB*, *plsC*, *cdsA*, *pssA*, *psd*, *pgsA* and *pgpA*, and two fragments with the fluorescent marker genes *eyfp-spinach* and *mCherry*. All genes are under the control of a T7 promoter, except *pgsA*, *pgpA* and *eyfp-spinach*, which are controlled by an SP6 promoter. The genes *ftsZ:mVenus* and *ftsA* are combined in an operon. Additionally, a fragment containing the oriR and oriL sequences with an internal PmeI restriction site is included, which allows linearization of the chromosome and replication with the φ29 DNA replication machinery. The remaining seven fragments contain sequences for amplification, screening and selection in *S. cerevisiae*: a yeast centromere and replication origin (CEN6/ARS4), and yeast screening and selection markers (*crtE*, *crtI*, *crtYB*, *mRuby2*, *hphNT1* and *URA3).* The CEN6/ARS4 fragment contains a landing pad and I-SceI restriction site for editing and linearization of the chromosome. The *crtI* fragment contains an additional yeast replication origin: ARS417. The *URA3* gene is combined on one fragment with a pCC1BAC backbone containing an antibiotic marker (*cat*) for amplification and selection in *E. coli*.

SHR overhangs of 60 bp (Supplementary data 3.10) were included in the PCR primers used to amplify all fragments (Supplementary data 3.6 and 3.7) to allow assembly in yeast by homologous recombination.

All assembly fragments were cleaned up by DpnI digestion and PCR purification or gel extraction, as described in the section “PCR and clean-up”. For each SynChr assembly, a mix of DNA fragments was prepared, containing 100 fmol of the yeast replication origin and *hphNT1* fragments and 150 fmol of all other fragments. After transformation, cells were plated on YPD + Hyg pates and incubated for three days at 30 °C. Plating on YPD + Hyg plates selects for transformants which have taken up both the *hphNT1* fragment, conferring resistance to hygromycin B, and the yeast replication origin fragment, placed on opposite locations on the SynChrs. Transformants were screened based on (i) β-carotene production, visible on plate as orange colonies, (ii) fluorescence measured by flow cytometry, and (iii) auxotrophy by streaking on SMD. The sequence of the SynChrs was determined by long-read sequencing of total DNA extracted from yeast.

Resulting strains harboring correctly and incorrectly assembled versions of MSG0.1, MSG0.2 and MSG1 were stocked for further characterization (Supplementary data 1.2).

### Assembly screening and sequencing

#### Fluorescence detection by flow cytometry

Yeast colonies containing assembled SynChrs were checked for mRuby2 (MSG0.1, MSG0.2 and MSG1 colonies)) and mTurquoise2 (MSG0.1 and MSG0.2 colonies) fluorescence by flow cytometry using the BD FACSAria II Cell Sorter with the BD FACSDiva software (BD Biosciences, Franklin Lakes, NJ, USA). Before every experiment, a cytometer Setup and Tracking (CS&T) cycle was run according to the manufacturer’s instructions. MSG0.1 and MSG0.2 colonies were resuspended in 500 µL Isoton II Diluent (Beckman Coulter, Woerden, The Netherlands). MSG1 colonies were resuspended in 500 µL synthetic medium without glucose and vitamins for direct measurement, or in 500 µL complete SMD followed by 4 h incubation at 30 °C at 200 rpm to wake up the cells and ensure active expression of the fluorescent markers before measurement. The 70 µm nozzle was used and at least 30,000 events per sample were recorded. A 561 nm laser with a 582/15 nm bandpass emission filter was used for detection of mRuby2, and a 445 nm excitation laser with a 525/50 nm bandpass filter for mTurquoise2 detection. Data analysis was performed with the Cytobank software (Beckman Coulter). The negative control strain CEN.PK113-7D and positive control strain IMF2 (Supplementary data 1.1) were used to determine the gates to distinguish fluorescent cells from non-fluorescent ones.

#### Auxotrophy detection by restreaking on SMD

The presence of the *URA3* fragment (MSG0.1, MSG0.2 and MSG1) and *LEU2* and *HIS3* fragments (MSG0.1 and MSG0.2) was determined by picking colonies from the YPD + Hyg transformation plate and streaking them on SMD. No growth on SMD indicated absence of at least one auxotrophic marker. In the absence of growth of MSG0.1 and MSG0.2 colonies on SMD, a second restreak was performed from the YPD + Hyg plate onto three SMD agar plates: (i) SMD + uracil + histidine, (ii) SMD + uracil + leucine, (iii) SMD + histidine + leucine. Plates were incubated for two days at 30 °C.

#### Long-read sequencing

To determine the sequence of MSG0.1 and MSG0.2 assembled in yeast, whole genome long-read sequencing was performed in-house using MinION technology (Oxford Nanopore Technologies, Oxford, UK). Genomic DNA was prepared and checked for purity and quantity using NanoDrop and Qubit as described above. DNA length and integrity were analyzed using a genomic DNA ScreenTape assay (Agilent) in the TapeStation 2200 (Agilent). Libraries of two to four barcoded samples were prepared with the ligation sequencing kit SQK-LSK109 and expansion kit EXP-NBD104 (Oxford Nanopore Technologies) according to the manufacturer’s recommendations. Three FLO-MIN106 (R9.4.1) flow cells were used to sequence the following strains (Supplementary data 1.2): (i) IMF49– IMF52, (ii) IMF53–IMF56, (iii) IMF59, IMF61, IMF62 and IMF64. For IMF53–IMF56, additional I-SceI digestion (Thermo Fisher Scientific) to linearize the SynChrs was performed according to the manufacturer’s instructions and the DNA was cleaned-up using AMPure XP beads (Beckman Coulter). Due to low sequencing read depth, strains IMF51 and IMF54 were additionally sequenced together on a FLO-MIN106 (R9.4.1) flow cell with adaptations to the library preparation to improve read length: fresh genomic DNA (five days old) was used, ligation and end repair times were extended to 10 min, adapter ligation was extended to 30 min, incubation time with magnetic beads at each step was extended to 10 min and elution from the beads was performed at 37 °C for 10 min. To increase output for IMF51 and IMF54, after loading one-third of the library and sequencing for 21 h, the flow cell was washed with kit EXP-WSH004 according to the manufacturer’s instructions and the other two-third of the library was loaded and sequenced for 45.5 h.

Guppy (GPU v6.3.4, Oxford Nanopore Technologies) was used for basecalling and demultiplexing. The resulting fastq files were filtered on length (> 200 bp for the first three sequencing runs, > 1 kb for the second run of IMF51 and IMF54) and mapped using minimap2 (parameters: *-ax map-ont*) ^70^ to a reference FASTA file containing the sequence of CENPK113-7D ^67^ concatenated with the designed SynChr sequence. The mapped reads were sorted and indexed using SAMTools ^73^ and visualized with the Integrated Genomics Viewer (IGV, version 2.11.9) ^72^. An annotation (.gff) file of the designed SynChr was created by exporting the annotations from SnapGene (version 7.0.2, GSL Biotech, San Diego, CA, USA) and using the R package labtools (version 0.1.0, function: *make_gff_from_snap*) ^68^ in R Statistical Software (version 4.1.2) ^69^. De novo assembly of the reads was performed using Flye version 2.7.1-b1673 (parameters: *--plasmids –nano-raw*) ^75^ or Canu version 2.2 (parameters: *genomeSize=12m, minReadLength=200, minOverlapLength=50, -nanopore-raw*) ^76^ and the assembled contig corresponding to the SynChr was identified with the MUMmer package ^77^, using NUCmer (parameter: *-maxmatch*) and delta-filter (parameter: *-q*). If necessary, a consensus SynChr sequence was assembled in SnapGene using information from the Flye and Canu assemblies and raw reads.

Strains harbouring MSG1 variants (Supplementary data 1.2) were sequenced by Plasmidsaurus using the “Big bacterial genome” service after total DNA extraction from yeast. The resulting .fna file containing all contigs was opened in SnapGene and the contig corresponding to the SynChr was identified by detection of common features. Some SynChr contigs were predicted to be linear by the analysis pipeline of Plasmidsaurus (involving de novo assembly of reads using Flye), which is expected to be incorrect. Whenever doubts were present about the accuracy of the consensus sequence, raw reads were analyzed as described for SynChrs isolated from *E. coli*. After transfer of the MSG1 variants from *S. cerevisiae* to *E. coli* and isolation from *E. coli*, the SynChrs were again sequenced by Plasmidsaurus, as described in the section “Plasmid and SynChr isolation from *E. coli* and verification”.

Raw reads of the total extracted DNA and consensus sequences of MSG variants are provided in Supplementary data 6. An overview of mutations in the MSG1 variants is provided in Supplementary data 4.5.

### SynChr stability

A single colony of IMF49 (MSG0.2) or IMF54 (MSG0.1) was used to inoculate three individual 50-mL centrifuge tubes containing 20 mL SMD or SMD supplemented with uracil and histidine. Cultures were grown overnight at 30 °C while shaking. For four days, the cells were kept at exponential phase by transferring to fresh medium every day. Cultures after 44 and 88 h were subjected to flow cytometry as described above. Additionally, 1 µL of culture was diluted in 999 µL of sterile water and 50 µL of this dilution was plated on SMD or SMD + uracil + histidine agar plates. Plates were incubated at 30 °C for two days.

### SynChr isolation from yeast for cell-free expression

Two isolations of MSG0.1 were performed (isolation 1 without replicates, isolation 2 with technical duplicates), which differed in starting yeast cell count. IMF54 was grown from glycerol stock in SMD until a final ODV (ODV = OD_660_ × Volume [mL]) of 1070 for isolation 1 and ODV of 9000 for isolation 2 (4500 per replicate). For isolation 1, cells were harvested by centrifugation at 4000 g for 5 min at 4 °C and the cell pellet was washed by resuspension in 4 mL TE buffer (10 mM Tris-HCl, 1 mM EDTA, pH 8.0). Washed cells were harvested by centrifugation at 4000 g for 5 min at 4 °C and resuspended in 4 mL Buffer Y1 (1 M sorbitol, 100 mM EDTA, 14 mM β-mercaptoethanol, pH 8.0, prepared in house or taken from Genomic DNA Buffer Set, Qiagen). To digest the cell walls and obtain spheroplasts, 12.5 mg Zymolyase 20T (AMSBIO, Alkmaar, The Netherlands), dissolved in 250 µL Milli-Q, was added and mixed with the cells through inversion. For isolation 2, the isolation of the SynChrs was performed in duplicate, starting from the same cell culture. Cells of both replicates were harvested together by centrifugation and washed in 33.6 mL TE buffer, harvested again and resuspended in 33.6 mL Buffer Y1, after which the cells were divided into two tubes to do the SynChr isolation in duplicate. To each duplicate, 52.6 mg Zymolyase 20T resuspended in 1051 µL Milli-Q was added.

The cells were incubated in a water bath at 30 °C for at least 1 h with regular mixing. Efficiency of spheroplasting was measured by diluting 5 µL of cells before and after spheroplasting in 995 µL demiwater and measuring the OD_660_. The low osmotic pressure of water will result in bursting of the spheroplasts. Once the OD_660_ was reduced by more than 80%, the spheroplasts were placed on ice and harvested by centrifugation at 4000 g for 15 min at 4 °C. The spheroplast pellet was resuspended in 1 mL 1 M sorbitol (isolation 1) or 2 mL 1 M sorbitol (isolation 2). MSG0.1 was isolated from the spheroplasts using the Qiagen Large-Construct Kit following the manufacturer’s instructions, starting from the addition of 19 mL (isolation 1) or 18 mL (isolation 2) Buffer P1 to have a total volume of 20 mL resuspended spheroplasts. The following adjustments were made to the manufacturer’s instructions: After elution in 15 mL Buffer QF, the eluate was stored overnight at 4 °C and the next morning, the DNA was precipitated with isopropanol in the presence of 15 µL 15 mg mL^−1^ GlycoBlue Coprecipitant (Invitrogen) and 1.2 mL 3M sodium acetate (pH 5.2, Sigma-Aldrich) to improve visibility of the DNA pellet. The final isolated MSG0.1 DNA was dissolved in 100 µL 10 mM Trizma hydrochloride solution (pH 8.5, Sigma-Aldrich).

MSG0.1 isolation from IMF54 was attempted with three other protocols, which failed to result in *yfp* expression in PURE system. The protocols are described below.

i. Cells with an ODV of 1912 (956 per replicate) were harvested together for both replicates through centrifugation at 4000 g for 5 min at 4 °C, resuspended in 20 mL 0.1 M Tris-HCl buffer (pH 8.0, Thermo Scientific, Thermo Fisher Scientific) and centrifuged again at 4000 g for 5 min at 4 °C. The cell pellet was resuspended in 2185 µL Tris-DTT solution (0.1 M Tris-HCl, 10 mM dithiothreitol (Roche, Sigma-Aldrich), prepared fresh before use) and divided into two tubes to do the SynChr isolation in duplicate. RNase A solution (20 mg mL^−1^, Sigma-Aldrich) was added into a final concentration of 100 µg mL^−1^ and the cells were incubated for 10 min at 30 °C in a shaking incubator at 200 rpm. Zymolyase 20T (13.65 mg) was dissolved in 136.5 µL Tris-DTT solution and added to the cells, followed by incubation for 15 min at 30 °C in a shaking incubator at 200 rpm. Efficiency of spheroplasting was determined as described above and the spheroplasts were placed on ice once the OD_660_ was reduced by more than 80%. Spheroplasts were not harvested through centrifugation, but directly divided over multiple tubes (250 µL per tube). The MSG0.1 plasmid was isolated using the GeneJET miniprep kit (Thermo Fisher Scientific) starting from the addition of 250 µL lysis buffer, following the manufacturer’s instructions with adaptations. After centrifugation of the cell debris and chromosomal DNA, the supernatant was not clear and was therefore transferred to a new tube and centrifuged again for 5 min at 15000 g, after which the resulting supernatant was loaded onto the column. At this step, the supernatants from all tubes were combined onto one column per replicate. MSG0.1 was eluted from the column by addition of 25 µL prewarmed Milli-Q (55 °C) and incubation for 10 min at RT before 2 min centrifugation.
ii. An adaptation of protocol (i) was performed, where lysis of spheroplasts by osmotic shock was avoided through changing the spheroplasting buffer from Tris-DTT solution to Buffer Y1. The MSG0.1 was isolated from the intact spheroplasts using the GeneJET miniprep kit, starting from addition of resuspension buffer. Cells were harvested at an ODV of 2140 (1070 per duplicate) by centrifugation at 4000 g for 5 min at 4 °C and the cell pellet was washed by resuspension in 8 mL TE buffer. Washed cells were harvested by centrifugation at 4000 g for 5 min at 4 °C and resuspended in 8 mL Buffer Y1. Zymolyase 20T (25 mg dissolved in 500 µL Milli-Q) was added and mixed with the cells through inversion. The cells were incubated in a water bath at 30 °C for 1 h 25 min with regular mixing. Spheroplasts were placed on ice and harvested by centrifugation at 4000 g for 10 min at 4 °C. The spheroplast pellet was resuspended in resuspension buffer from the GeneJET miniprep kit to have a total volume of 2.5 mL and was divided over 10 tubes (250 µL per tube, 5 tubes per duplicate). The MSG0.1 was isolated in duplicate from the resuspended spheroplasts using the GeneJET miniprep kit, following manufacturer’s instructions from the addition of 250 µL lysis buffer to each tube. After centrifugation to pellet the cell debris and chromosomal DNA, each supernatant was transferred to a new tube and centrifuged again for 5 min at 15000 g. For each replicate, five supernatants were combined onto one column. MSG0.1 was eluted from the column by addition of 25 µL prewarmed Milli-Q (55 °C) and incubation for 10 min at RT before 2 min centrifugation.
iii. The published protocol by Noskov and colleagues ^78^ was attempted twice with technical duplicates.

### Cell-free gene expression

In vitro transcription-translation of MSG0.1 isolated from yeast and of MSG1 isolated from EPI300 *E. coli* was performed using PURE*frex*2.0 (GeneFrontier Corporation, Kashiwa, Japan), following the manufacturer’s instructions for storage and handling. Cell-free expression was performed in bulk (test tube) reactions (MSG0.1 and MSG1) and in liposomes (MSG1 only).

Bulk reactions were prepared in standard PURE*frex* 2.0, containing T7 RNAP for transcription, or in custom PURE*frex* 2.0 without T7 RNAP and with addition of SP6 RNAP (Promega). PURE reactions with T7 RNAP were prepared by mixing 5 µL solution I (buffer), 0.5 µL solution II (enzymes), 1 µL solution III (ribosomes), 5 U SUPERase·In (Thermo Fisher Scientific), DNA template and Milli-Q in a final volume of 10 µL. PURE reactions with SP6 RNAP were prepared by mixing 5 µL solution I, 0.5 µL solution II ΔT7 RNAP, 20 U SP6 RNAP, 1 µL solution III, 5 U SUPERase·In, DNA template and Milli-Q in a final volume of 10 µL.

### Cell-free expression of MSG0.1 isolated from yeast

#### DNA templates

Positive control templates for *yfp* expression were plasmids G76 (isolation 1) and pUD1251 (isolation 2), isolated from *E. coli* (Supplementary data 2.5), and PCR products of the *yfp* cassette (isolation 2 only), amplified using Phusion polymerase and primer pair 19274/19277 (Supplementary data 3.9) from the MSG0.1 isolated from IMF54. Plasmid G28 (Supplementary data 2.5) was used to determine PURE expression inhibition by the MSG0.1 sample (isolation 1 and 2). The *mCherry* gene encoded on G28 was expressed in PURE system in the presence of MSG0.1 sample, water (positive control) or non-coding plasmid DNA (positive control). Plasmid G28 (0.5 µL, resulting in 1 nM final plasmid concentration), diluted in 2.75 µL MSG0.1 isolated from IMF54, 2.75 µL water or 2.75 µL 30 ng µL^−1^ non-coding plasmid pUDC192, was added to the reaction mixture.

Reactions were assembled in PCR tubes and transferred to a black 384-well flat µClear bottom microplate (Greiner Bio-One, Kremsmünster, Austria), which was sealed with a highly transparent film (Sarstedt, Nümbrecht, Germany).

#### Spectrofluorometry

The plate was incubated at 37 °C for 16 h and fluorescence was measured at end-point in triplicates in an Infinite 200 Pro plate reader (Tecan, Männedorf, Switzerland) using bottom and top measurements (isolation 1) or a CLARIOstar plate reader (BMG LABTECH, Ortenberg, Germany) using bottom measurement (isolation 2). YFP fluorescence was measured with excitation at 488 nm (Infinite 200 Pro) or 497 nm with 15 nm bandwidth (CLARIOstar) and emission at 527 nm (Infinite 200 Pro) or 540 nm with 20 nm bandwidth (CLARIOstar). For mCherry, fluorescence was detected with excitation at 570 nm (Infinite 200 Pro) or 559 nm with 20 nm bandwidth (CLARIOstar) and emission at 620 nm (Infinite 200 Pro) or 630 nm with 40 nm bandwidth (CLARIOstar). For measurements in the Infinite 200 Pro, a gain of 80% was used. For measurements in the CLARIOstar, the focus and gain were adjusted to a well containing a PURE*frex*2.0 reaction solution with 70 pM pUD1251 (YFP) or 1 nM G28 (mCherry), that was pre-incubated for five hours at 37 °C in a thermocycler. The gains used in the CLARIOstar were 1852 (YFP) and 2090 (mCherry) and the focus height 3.1 mm.

### Cell-free expression of MSG1 isolated from *E. coli*

#### DNA templates

DNA templates were measured by Qubit and added in a final concentration of 1 nM (MSG1 variants and control plasmids). Plasmids G28, G200, G276, G396, G435 and G613 (Supplementary data 2.6) were used as control plasmids for expression of individual modules. All reactions were performed in biological triplicate.

Reactions were assembled in PCR tubes and incubated at 37 °C for 16 h, either in PCR tubes in a thermocycler (samples containing control plasmids G276, G396, G435 or G613) or after transfer to a 384-well plate in a spectrofluorometer (samples containing SynChrs or control plasmids G28, G200 or G613) as described below. After incubation, samples were flash-frozen in liquid nitrogen and stored at −80 °C.

#### Spectrofluorometry

Bulk reactions containing SynChrs or control plasmids G28, G200 or G613 were transferred to a black 384-well flat µClear bottom microplate (Greiner Bio-One, Kremsmünster, Austria), which was sealed with a highly transparent film (Sarstedt, Nümbrecht, Germany). The plate was incubated at 37 °C for 16 h in a Synergy H1 Multi-Mode Microplate Reader (Agilent) and bottom fluorescence was measured every 5 min using an excitation bandwidth of 500/20 nm and emission bandwidth of 539/20 nm for detection of eYFP and mVenus fluorescence, and with excitation at 579/20 nm and emission at 616/20 nm for mCherry detection. The read height was set to 9.5 mm and a gain of 100 was used for mCherry and 50 for eYFP and mVenus.

Fluorescence data were analyzed using a custom Python script that automated processing and plotting of kinetic curves. Gene expression kinetics were modeled using a sigmoid function based on ^45^ to fit the experimental data, allowing extraction of the maximum apparent translation rate (RFU h^−1^).

Maximum fluorescence (RFU) was determined using a sliding window approach, identifying the 100-min time period with the highest average fluorescence. The maximum fluorescence was then calculated by averaging the fluorescence values of the 20 data points within this window.

### Liquid chromatography mass spectrometry

#### Sample preparation for label-free proteomics analysis

Bulk reactions containing SynChrs (T7 RNAP and SP6 RNAP samples) or control plasmids G276, G396, G435 or G613 (T7 RNAP samples only) were analyzed by liquid chromatography mass spectrometry (LC-MS). Ten microliter of protein samples were processed for trypsin digestion by addition of ten microliter of trypsin (500 ng) in ammonium bicarbonate (100 mM) to each sample for overnight digestion. The reaction was stopped by adding 1.5 µL 1% trifluoroacetic acid (TFA).

#### NanoLC-MS/MS analysis of proteins

Tryptic peptides were analyzed by nano-LC coupled to tandem MS, using an UltiMate 3000 system (NCS-3500RS Nano/Cap System, Thermo Fisher Scientific) coupled to an Orbitrap Q Exactive Plus mass spectrometer (Thermo Fisher Scientific). Five microliter of sample was injected on a C18 precolumn (300 µm inner diameter × 5 mm, Thermo Fisher Scientific) in a solution consisting of 2% acetonitrile and 0.05% TFA, at a flow rate of 20 µL min^−1^. After 5 min of desalting, the precolumn was switched online with the analytical C18 column (75 μm inner diameter × 50 cm; in-house packed with Reprosil C18) equilibrated in 95% solvent A (5% acetonitrile, 0.2% formic acid) and 5% solvent B (80% acetonitrile, 0.2% formic acid). Peptides were eluted using a 10–50% gradient of solvent B over 105 min at a flow rate of 300 nL min^−1^. The mass spectrometer was operated in data-dependent acquisition mode with the Xcalibur software. MS survey scans were acquired with a resolution of 70,000 and an AGC target of 3×10^6^. The ten most intense ions were selected for fragmentation by high-energy collision-induced dissociation, and the resulting fragments were analyzed at a resolution of 17,500 using an AGC target of 1×10^5^ and a maximum fill time of 50 ms. Dynamic exclusion was used within 30 s to prevent repetitive selection of the same peptide.

#### Bioinformatics analysis of MS raw files

Raw MS files were processed with the Mascot software (version 2.7.0) for database search and Proline ^79^ for label-free quantitative analysis (version 2.1.2). Data were searched against *E. coli* entries of the UniProtKB protein database release Swiss-Prot 2019_11 (23,135 entries) and homemade database (built with FASTA sequences of expected proteins mVenus from *Aequorea victoria*, DNAP and TP from *Bacillus* phage φ29 and mCherry from *Anaplasma marginale*). Oxidation of methionine was set as a variable modification. Specificity of trypsin/P digestion was set for cleavage after K or R, and two missed trypsin cleavage sites were allowed. The mass tolerance was set to 10 ppm for the precursor and to 20 mmu in tandem MS mode. Minimum peptide length was set to 7 amino acids, and identification results were further validated in Proline by the target decoy approach using a reverse database at both a PSM and protein false-discovery rate of 1%. For label-free relative quantification of proteins across biological replicates and conditions, cross-assignment of peptide ion peaks was enabled within each group with a match time window of 1 min, after alignment of the runs with a tolerance of ± 600 s. Raw abundance values were visualized as bar plots.

A two-way ANOVA with post-hoc Tukey testing was performed on log_2_-transformed raw abundance values to analyze variation across DNA templates and proteins.

A matrix of abundance ratios was generated for each pair of runs using Median Ratio Fitting, based on ion abundances for each protein. For each pairwise comparison, the median of the ion abundance ratios was then calculated and used to represent the protein ratio between these two runs. A least-squares regression was performed to estimate the relative abundance of the protein across all runs in the dataset. Finally, these abundance values were rescaled to the total sum of the ion abundances across runs.

To assess differences in protein abundance between biological groups, a two-tailed Student’s *t*-test (equal variances) was performed on log_2_-transformed abundance values. Significance level was set at *p* = 0.1 and ratios were considered relevant if higher than ± 2. Results were visualized using volcano plots.

Lastly, principal component analysis (PCA) was conducted on the rescaled abundance values to identify potential outliers.

LC-MS data underlying Figure 5 and Supplementary Figure 12 are provided in Source data 1.

### Preparation of reactions with custom PURE*frex*

Custom PURE*frex* 2.0 was used for preparation of 20 µL PURE reactions in 1.5 mL tubes. MSG1.1 isolated from EPI300 *E. coli* (pUDF006) was added at a final concentration of 0.1 nM. A positive control reaction was prepared using a mixture of 0.1 nM linear *yfp* template (obtained by PCR-amplification of the *yfp* expression cassette from G76 using primers M13F and M13R (Supplementary data 3.9)) and 0.1 nM of plasmid G28, encoding *mCherry* (Supplementary data 2.6).

### Liposome preparation

Lipid-coated beads were prepared as described in references ^38,45^ with minor modifications. Glass beads (0.6 g, Sigma-Aldrich) were coated with 2 mg of 50 mol% DOPC, 36 mol% DOPE, 12 mol% DOPG, 2 mol% 18:1 cardiolipin, 0.05% (by mass) DOPE-Cy5, and 1% (by mass) DSPE-PEG-biotin. All lipids were from Avanti Polar Lipids.

Lipid-coated beads were desiccated for at least 30 min before use and 10–15 mg of beads was added to each PURE sample. The samples were gently rotated in an automatic tube rotator (VWR) for 1 h at 4 °C to allow liposome swelling, followed by four cycles of freezing by dipping in liquid nitrogen for 10 s and thawing for 5–10 min on ice. About 12 µL of the upper solution in each liposome sample was transferred to a PCR tube containing 0.5 µL DNase I (2 U µL^−1^, New England Biolabs), using a cut pipette tip, and the solution was gently pipetted up and down twice for mixing. Samples were incubated in a thermocycler for 14 h at 37 °C.

### Confocal microscopy

The wells of a black 384-well flat µClear bottom microplate were functionalized with 50 μL of 1 mg mL^−1^ BSA (Sigma-Aldrich). The plate was closed with a reusable cover and incubated overnight at 4 °C. The next morning, the BSA solution was pipetted out of the wells and the wells were washed twice with 10 μL Milli-Q, after which 10 μL homemade PURE buffer (180 mM potassium glutamate, 14 mM magnesium acetate, 20 mM HEPES-KOH at a pH of 7.6) was added. Liposome samples were diluted by mixing 5 µL homemade PURE buffer with 7 µL liposome sample using a cut pipette tip and slowly pipetting up and down three times. All PURE buffer was pipetted out of the wells and immediately 12 µL of diluted liposome sample was added. Liposomes were imaged after 20 min incubation at room temperature, allowing the liposomes to sink to the bottom of the wells. The Leica SP8 inverted confocal microscope equipped with a 63× objective and appropriate laser lines was used.

Confocal images were processed using Fiji (ImageJ version 1.53c) ^80,81^. Brightness was adjusted for each channel separately to improve visibility of the fluorescence signal and the three channels were combined into a composite image.

### DNA replication

DNA replication of MSG1 linearized by PmeI was performed using purified φ29 DNA replication proteins in an adjusted PURE*frex* 2.0 reaction, supplemented with the required substrates and cofactors for DNA replication. The concentration of tRNAs and NTPs was changed in PURE*frex* 2.0 (see the modified composition below) to allow DNA replication. Full-length replication was verified by agarose gel electrophoresis.

#### Preparation linear MSG1

SynChr MSG1.1 isolated from EPI300 *E. coli* (pUDF006) was linearized by PmeI (Thermo Fisher Scientific) following the manufacturer’s instructions, at a final DNA concentration of 100 ng µL^−1^. Linearization was verified by comparison of the linearized sample with the circular template on a 0.6% agarose gel in 1× TAE buffer, with the Quick-Load 1 kb Extend ladder as reference. Linear MSG1.1 was not purified from the restriction reaction. Final DNA concentration was quantified by Qubit.

#### Purified φ29 DNA replication proteins

Purified φ29 DNA replication proteins were produced as described in ^38^. Stock concentrations and storage buffers were: DNAP (420 ng μL^−1^ in 25 mM Tris-HCl (pH 7.5), 0.5 M NaCl, 0.5 mM EDTA, 3.5 mM β-mercaptoethanol (BME), 0.025% Tween 20, 50% glycerol), TP (320 ng μL^−1^ in 50 mM Tris-HCl (pH 7.5), 0.5 M NaCl, 1 mM EDTA, 7 mM BME, 50% glycerol), SSB (10 mg mL^−1^ in 50 mM Tris-HCl (pH 7.5), 60 mM (NH_4_)_2_SO_4_, 1 mM EDTA, 7 mM BME, 50% glycerol), DSB (10 mg mL^−1^ in 50 mM Tris-HCl (pH 7.5), 450 mM (NH_4_)_2_SO_4_, 1 mM EDTA, 7 mM BME, 50% glycerol). The proteins were aliquoted and stored at −80 °C.

#### Reaction conditions

A DNA replication reaction was prepared without ribosomes (to prevent expression of MSG1- encoded DNA replication proteins) by mixing 5 µL custom PURE*frex* 2.0 solution I (ΔtRNAs, ΔNTPs), 0.33× dNTPs, 0.33× tRNAs, 1 µL PURE*frex* 2.0 solution II, 12 U SUPERase·In, 20 mM (NH_4_)_2_SO_4_, 3 ng µL^−1^ DNAP, 3 ng µL^−1^ TP, 375 ng µL^−1^ SSB, 105 ng µL^−1^ DSB, 300 µM dCTP, 300 µM dGTP, 300 µM dATP, 180 µM dTTP, 120 µM fluorescein-12-dUTP (Thermo Fisher Scientific), 100 ng DNA template and Milli-Q in a final volume of 20 µL. Another reaction was prepared in duplicate in the same way, but with the addition of 2 µL PURE*frex* 2.0 solution III (containing ribosomes). Of each reaction, 10 µL was taken and stored in a PCR tube at −20 °C (t = 0 h samples). The remaining 10 µL was incubated in a PCR tube at 30 °C in a thermocycler for 16 h (t = 16 h samples).

#### Visualization on agarose gel

All 10 µL samples were incubated with 0.2 mg mL^−1^ RNase A (Promega) at 30 °C in a thermocycler for 1.5 h to remove RNA. The reactions were quenched by addition of 6 µL STOP solution (30 mM EDTA, 0.3% SDS) and proteins were removed using 1 mg mL^−1^ Proteinase K (Thermo Fisher Scientific) and 1 h incubation at 50 °C in a thermocycler. The samples were stored in the dark at 4 °C until visualization on gel.

The samples were analyzed by agarose gel electrophoresis on a 0.6% agarose gel without DNA stain in 1× TAE buffer, with the Quick-Load 1 kb Extend ladder as reference. The gel was run for 90 min at 53 V and visualized on a fluorescence gel imager (Typhoon, Amersham Biosciences) using a 488 nm excitation laser with 525/20 nm bandpass emission filter to detect fluorescein-labeled DNA. Post-staining was done in GelRed Nucleic Acid Stain (Millipore, Merck, Burlington, MA, USA), according to the manufacturer’s instructions, and the gel was washed twice for 5 min in Milli-Q. Total DNA was visualized on the fluorescence gel imager using a 532 nm excitation laser with a 570/20 nm bandpass filter to detect GelRed.

### Statistics and reproducibility

The number of replicates is specified in the figure legends. A representative image of gel is displayed in Supplementary Fig. 13; images from a repeated measurement are reported in Source data. Statistical tests for mass spectrometry data analysis have been performed as described in Methods.

## Data and code availability

Data are available in the main manuscript and the Supplementary Information. Source data are provided with this paper. LC-MS data underlying Figure 5 and uncropped gel images of Supplementary Figure 13 are available in Source Data. The code used to extract gene expression kinetic parameters from fluorescence measurements is available on the Github repository (https://github.com/DanelonLab/Kinetic-Analysis-of-PURE-System-Fluorescence-Data). The mass spectrometry proteomics data have been deposited to the ProteomeXchange Consortium via the PRIDE partner repository with the dataset identifier PXD069407.

## Acknowledgements

We thank the following people for constructing and sharing plasmids: Quinte Smitskamp (pUDE1217), David Foschepoth (G131), Nicole Bennis (pUDE1110), Matic Kostanjšek (pUD1226), Jasmijn Hassing (pUDE1039), Kristel Doets from Wageningen University and Research (pCC1BAC- lacZα), Ramon van de Valk and Liedewij Laan (pWKD014), Elynor Moore (pUD1394), Sophie van der Horst (G162, G197 and G200) and Federico Ramírez Gómez (G607 and G613).

We thank GeneFrontier for supporting our research, Álvaro Eseverri and Emma Barahona (Centre for Plant Biotechnology and Genomics, Spain) for providing genomic DNA of *Arabidopsis thaliana*, Marijke Luttik for technical assistance with FACS analysis and the development of the SynChr isolation protocol, Bastiaan van Dijk for help with developing SynChr isolation protocol, Clara Carqueija Cardoso for help with library preparation for long-read sequencing, Marcel van den Broek for help with sequencing data analysis, Yannick Bernard-Lapeyre for calculation of the maximum fluorescence and apparent translation rates from fluorescence data, and Marijn van den Brink for help with liposome preparation, confocal microscopy and flow cytometry. Confocal imaging was performed at the Light Imaging Toulouse CBI (LITC) facility.

Images in Figures 2, 4 and 6 and Supplementary Figure 7 were (partially) created with BioRender.com.

This work was financially supported by The Netherlands Organization for Scientific Research (NWO/OCW) via the “BaSyC – Building a Synthetic Cell” Gravitation Grant (024.003.019) and by Agence Nationale de la Recherche (ANR-22-CPJ2-0091-01). The project was also supported by the Région Occitanie, European funds (“Fonds Européens de Développement Régional”, FEDER), Toulouse Métropole, and by the French Ministry of Research with the “Investissement d’Avenir Infrastructures Nationales en Biologie et Santé” program (ProFI, Proteomics French Infrastructure project, ANR-10-INBS-08 & ANR-24-INBS-0015).

## Author contributions

P.D.L. and C.D. conceptualized and supervised the research and acquired funding. All authors designed the experiments; C.C., L.S.H. and E.Z. performed the experiments, analyzed the data and designed the figures; E.B. and A.S. performed the LC-MS experiments and analyzed the data; C.C., P.D.L. and C.D. wrote the paper, with input from all authors.

## Competing interests

The authors declare no competing interests.

## Supplementary information

Supplementary Data, Notes and Figures are available online. Supplementary data include four files in Excel format with information about strains, plasmids, linear DNA templates, primers, SHRs, and SynChrs used and constructed in this study (Supplementary data 1–4). The designed maps of MSG variants in GenBank format (Supplementary data 5), sequencing data after total DNA isolation from *S. cerevisiae* strains (raw Nanopore sequencing reads in FASTQ format and consensus SynChr sequences in GenBank format, Supplementary data 6) and sequencing data of MSG1 variants after isolation from *E. coli* (raw Nanopore sequencing reads in FASTQ format and consensus SynChr sequences in GenBank format, Supplementary data 7) are available in the 4TU.ResearchData repository with DOI 10.4121/4eb28eac-b94c-4e97-9db6-4cd3af73fad2.v1.

## Notes

https://github.com/DanelonLab/Kinetic-Analysis-of-PURE-System-Fluorescence-Data

https://data.4tu.nl/datasets/4eb28eac-b94c-4e97-9db6-4cd3af73fad2/1

### Competing Interest Statement

The authors have declared no competing interest.

