## Supplementary Information for "Assembly and cell-free expression of a partial genome for the synthetic cell"

#### Content

Pages 2 – 5 Supplementary Notes

Page 2 Supplementary Note 1: Assembly screening of MSG0.1 and MSG0.2

Page 2 Supplementary Note 2: Nanopore sequencing results of MSG0.1 and MSG0.2

Page 3 Supplementary Note 3: MSG0.1 and MSG0.2 are stable during growth

Page 4 Supplementary Note 4: MSG0.1 isolated from *S. cerevisiae* can be directly expressed in PURE system

Pages 6 – 20 Supplementary Figures 1–13

Page 21 Supplementary References

### Supplementary Notes

#### Supplementary Note 1: Assembly screening of MSG0.1 and MSG0.2

We obtained a total of 53 MSG0.1 and 29 MSG0.2 colonies for screening step 1 (Supplementary Figure 1a). Visual inspection for the presence of carotenoid marker genes (step 2) showed that for MSG0.1, 25% of the colonies (13 out of 53 colonies) were orange, while 30% were white (16 colonies) and 45% (24 colonies) displayed mixed colors (Supplementary Figure 1). For the MSG0.2, 90% (26 out of 29) colonies were orange, 7% (2 colonies) were white, and a single colony (3%) displayed a mixed phenotype (Supplementary Figure 1). The results confirm the expected higher assembly efficiency in the absence of PURE repeats.

Mixed phenotypes reveal the presence of mixed populations that might result from different events. Upon plating after transformation, several adjacent cells could form a single colony. Alternatively, colonies could result from single cells that harbor multiple SynChrs with different configurations that segregate into separate daughter cells during division. Finally, mixed population could result from SynChr instability and recombination upon cell propagation. The possible occurrence of SynChr instability was investigated as described in Supplementary Note 3. Irrespective of the underlying events, the presence of PURE repeats and resulting undesired recombination events increased the frequency of colonies with mixed phenotype compared to MSG0.2 control colonies.

Through the following screening steps, five out of the 13 orange MSG0.1 colonies (representing 9% of total colony count in step 1) were positively screened for both fluorescent and auxotrophic markers (Supplementary Figure 1b), suggesting correct assemblies. For MSG0.2, 25 out of the 26 orange colonies carried the two fluorescent and the three auxotrophic markers (representing 86% of the total colonies from step 1) (Supplementary Figure 1b). Data from individual transformations are reported in Supplementary Figure 2.

Finally, we explored the potential of the 2μ replication origin by performing a 20-fragment assembly for both MSG0.1$^{2\mu}$ and MSG0.2$^{2\mu}$. For MSG0.2$^{2\mu}$, only one of the resulting six colonies was orange (Supplementary Figure 3a, c). Further screening of the orange colony for the presence of *mRuby2* and *mTurquoise2* fragments revealed a mixture of phenotypes with and without fluorescence and the colony was therefore classified as incorrect (Supplementary Figure 3b). For MSG0.1$^{2\mu}$, none of the 12 colonies displayed an orange phenotype (Supplementary Figure 3a, c). Due to the low assembly efficiency revealed by this preliminary data, further work on the 2μ version was discontinued.

#### Supplementary Note 2: Nanopore sequencing results of MSG0.1 and MSG0.2

In-depth inspection of the sequences showed that the assembled MSGs in strains IMF51 and IMF54 were near-perfect replicas of the in silico design (differing only by five point mutations for IMF51 and seven point mutations for IMF54). The total expected size of MSG0.1 was 69 kb, however, due to variations in DNA fragment size between the *A. thaliana* reference sequence used for the design and the final size in the template plasmids, the size of the constructed MSG0.1 was 67 kb. The assembled SynChrs of IMF50 and IMF52 showed some minor deviations that were most likely not caused by the presence of PURE repeats (Supplementary Figure 4). IMF50 missed a 1,992-bp sequence in the middle of the fragment with CEN6/ARS4, leading to the absence of ARS4, bla and ColE1 (Supplementary Figure 5). The missing sequence does not contain similarity with other parts of MSG0.1, indicating that unexpected homologous recombination events were not responsible for misassembly. IMF52 harbored an insertion of a 1,780-bp sequence in the terminator region downstream of PF9, corresponding to a plasmid backbone used as template during PCR, which suggests that this backbone was a contaminant in the transformation mix.

To explore if PURE repeats were involved in misassemblies of MSG0.1, six strains that did not pass the screening due to the absence of at least one marker were sequenced (strains IMF55, IMF56, IMF59, IMF61, IMF62 and IMF64, Supplementary Figure 6, Supplementary data 1.2). Based on the phenotype observed during the different screening stages, configurations could be predicted for the misassembled plasmids (Supplementary Figure 6). For white colonies, no prediction could be made as to which of the three markers for β-carotenoid were missing. Four out of the six strains (IMF55, IMF56, IMF561 and IMF62) showed the predicted configuration. Recombination had occurred between either promoter or terminator regions. The strain IMF55, similarly to IMF50, missed part of the CEN6/ARS4_*bla*_ColE1 fragment (2,076 bp, Supplementary Figure 5). For IMF59, a duplication of PF10 and the *HIS3* fragment had occurred, which was inserted in the promoter region upstream PF9 (Supplementary Figure 6), likely due to homologous recombination between PURE repeats. For IMF64, no consensus sequence could be determined because of a low depth of sequencing read. These results show that for most sequenced SynChrs, PURE repeats were responsible for misassemblies. Notably, recombination between SHRs is more frequent than between PURE repeats in the selected strains. Moreover, our screening pipeline successfully predicts SynChr configurations, even though it does not detect duplications or insertions, as expected.

A single strain harboring control plasmid MSG0.2 (IMF49) with all expected marker fragments was sequenced. Based on previous studies reporting a relatively high assembly efficiency (36%) for 44 fragments [1], an even higher efficiency was expected for MSG0.2 from 20 fragments, eliminating the need for sequencing multiple strains. The sequencing results confirmed the presence of all markers and DNA fragments with the expected configuration (Supplementary Figure 4). However, an 89-bp insert was present after the first SHR of CF4 and, similarly to what was observed for IMF50 and IMF55, part of the CEN6/ARS4_*bla*_ColE1 fragment (1,891 bp) showed approximately half the sequencing depth of the rest of control MSG0.2 (Supplementary Figure 5).

#### Supplementary Note 3: MSG0.1 and MSG0.2 are stable during growth

An important requirement for *S. cerevisiae* to become a genome foundry for PURE-based synthetic cells is the stability of the assembled plasmids in yeast cells during growth. The high incidence of colonies with mixed phenotypes (45%) for the MSG0.1 assembly might be caused by post-assembly homologous recombination events between PURE repeats. As HDR is primarily active during cell division (S and G2 phases of the mitotic cell cycle) [2], the risk of recombination events caused by the presence of repeats might increase during cell propagation. Many generations are required for completion of the workflow, from plating of the transformation mix, to screening, storing and finally propagating for SynChr extraction. The stability of the assembled plasmids during propagation was therefore explored. Strain IMF54 harboring a correctly assembled MSG0.1 and strain IMF49 carrying MSG0.2 were grown in liquid medium, and subjected to four serial transfers (Supplementary Figure

7a). Two different media were used to exert different levels of selection pressure for the maintenance of integer plasmids. Growth on SMD without supplement selected for the presence of all three auxotrophic markers *HIS3*, *LEU2* and *URA3*, while survival on SMD supplemented with uracil and histidine only required the presence of a single marker (*LEU2*, Figure 1). The integrity of the synthetic chromosomes was verified by measuring mRuby2 and mTurquoise2 fluorescence by flow cytometry, and by plating combined with visual inspection for β-carotene production after 44 and 88 hours of growth in culture (ca. 16 and 32 generations, respectively) (Supplementary Figure 7a). For both strains (IMF49 and IMF54), the percentage of cells with mTurquoise2 fluorescence remained above 98% after 88 hours, in both selection conditions (Supplementary Figure 7b). The percentage of cells showing fluorescence was not significantly affected by the difference in selection pressure (two-way ANOVA with Post-Hoc Tukey-Kramer, $p > 0.005$). Similar results were obtained after 44 hours and for mRuby2 (Supplementary Figure 7a-c). On plates, only 0 to 3 white colonies were observed on a total of more than 500 colonies per plate (Supplementary Figure 7c-e). Overall, our results indicate that SynChrs are stable during propagation in yeast and that the presence of PURE repeats does not hamper stability. We therefore speculate that the population heterogeneity shown by the mixed colors of 45% of the transformants may have occurred earlier during SynChr assembly or may be due to the fact that two different colonies converged on the same physical location on the plate.

#### Supplementary Note 4: MSG0.1 isolated from *S. cerevisiae* can be directly expressed in PURE system

Multiple protocols were attempted for SynChr isolation from *S. cerevisiae*, of which only one resulted in *yfp* expression from MSG0.1 in PURE system. All tested protocols are described in Methods under "SynChr isolation from yeast for cell-free expression". We noticed that careful yeast spheroplasting to avoid osmotic lysis, followed by column purification using a kit designed for large-construct isolation, was essential to extract sufficient amounts of MSG0.1. Two isolations were performed using the final protocol, which differed in starting yeast cell count (ca. four times more for the second isolation). Total concentration of DNA was measured by Qubit (Supplementary Figure 8a, isolation 2 only). The concentration of MSG0.1 was determined by qPCR using primers targeting specific regions on the SynChr (Supplementary Figure 8b).

For isolation 2, about 10% of total DNA isolated from strain IMF54 corresponded to MSG0.1 (Supplementary Figure 8a, b). Approximately 300 ng of MSG0.1 DNA could be extracted, which was sufficient for assaying expression of the encoded *yfp* gene in PURE system. Starting from about 20 pM of MSG0.1 template, clear production of YFP was detected by spectrofluorometry after 16h of incubation, compared to the negative control containing water instead of DNA (Supplementary Figure 8c). We also performed a series of control reactions using two reference templates: a plasmid isolated from *E. coli* coding for the YFP reporter protein (added at various concentrations, one of which similar to the MSG0.1 concentration) and a *yfp* transcriptional unit that was PCR-amplified from MSG0.1 (3.25 pM final concentration). Both control reactions revealed a higher relative yield of synthesized YFP per DNA template than with MSG0.1. As the lower yield may result from inhibitory effects from the DNA extract (e.g., the presence of native yeast DNA), we mixed a plasmid encoding mCherry with the MSG0.1 extract and measured its fluorescence after 16h (Supplementary Figure 8d). No significant drop of mCherry signal could be measured as compared to the control condition, where the MSG0.1 solution was substituted with water or a non-coding plasmid isolated from *E. coli*, indicating that the MSG0.1 extract did not visibly inhibit expression in PURE system. Most likely, the presence of the nine other transcription units present in MSG0.1 (vs. a single transcription unit in the reference templates) may lower the concentration of free T7 RNA polymerase, thus reducing the amounts of available resources for *yfp* transcription. Additionally, the lack of accuracy in determining the concentration of MSG0.1 could impact the results.

### Supplementary Figures

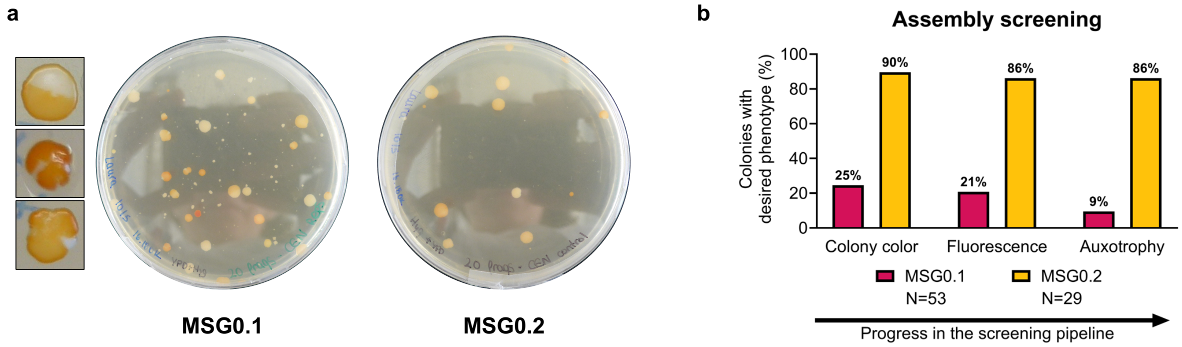


**Supplementary Figure 1*: Screening for SynChrs with correct assembly.* a)** Example of plates with YPD and hygromycin after transformation. MSG0.1 (left) and MSG0.2 (right). Zoom-in images show colonies with mixed phenotype regarding the presence of carotenoid genes. **b)** Percentage of colonies with desired phenotype at each screening step, compared to step 1. Colonies that grew on YPD with hygromycin plates were classified as positive when displaying an orange color, when showing fluorescence for both mTurquoise2 and mRuby2, and when growth on SMD was visible. Aggregated data from three independent transformations are displayed.

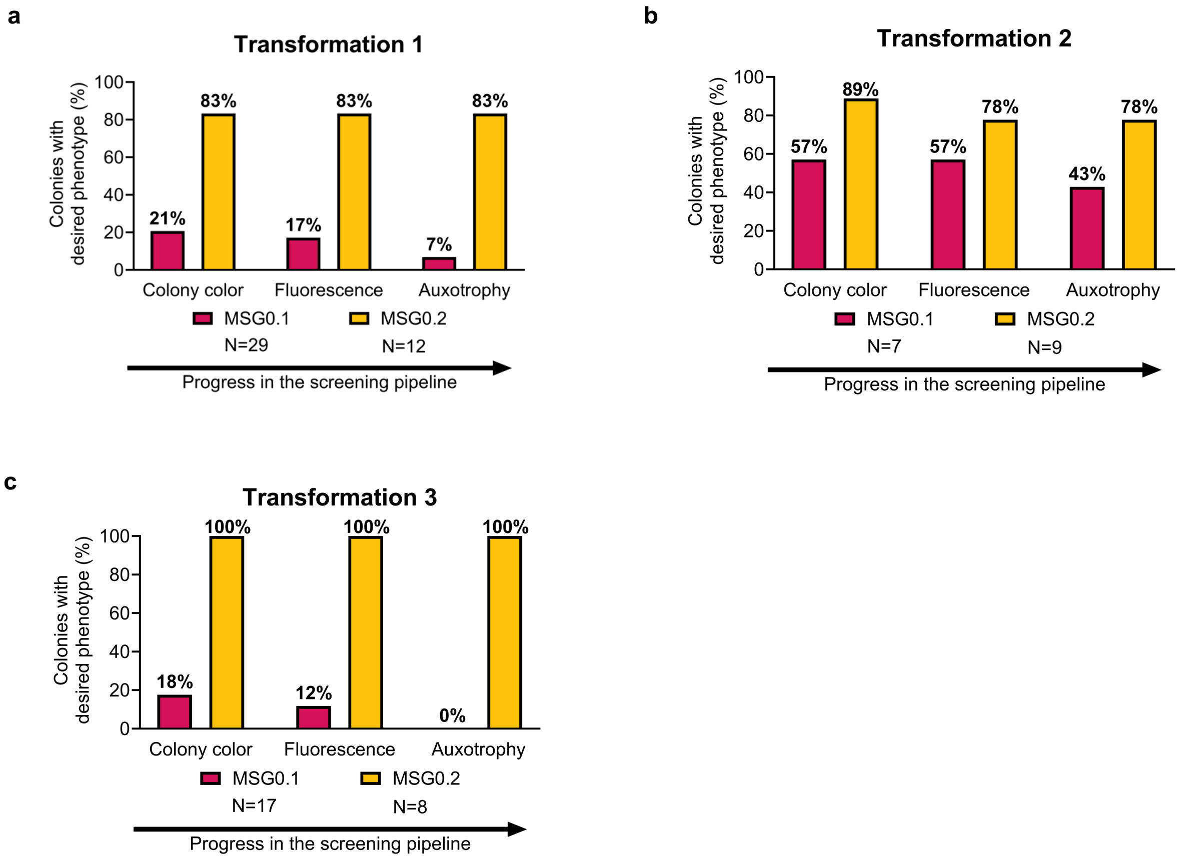


**Supplementary Figure 2: Percentage of colonies with desired phenotype for three independent transformations.** Colonies that grew on YPD with hygromycin plates were considered as correctly assembled when displaying an orange color, when showing fluorescence for both mTurquoise2 and mRuby2, and when growth on SMD was visible. **a)** Transformation 1. **b)** Transformation 2. **c)** Transformation 3.

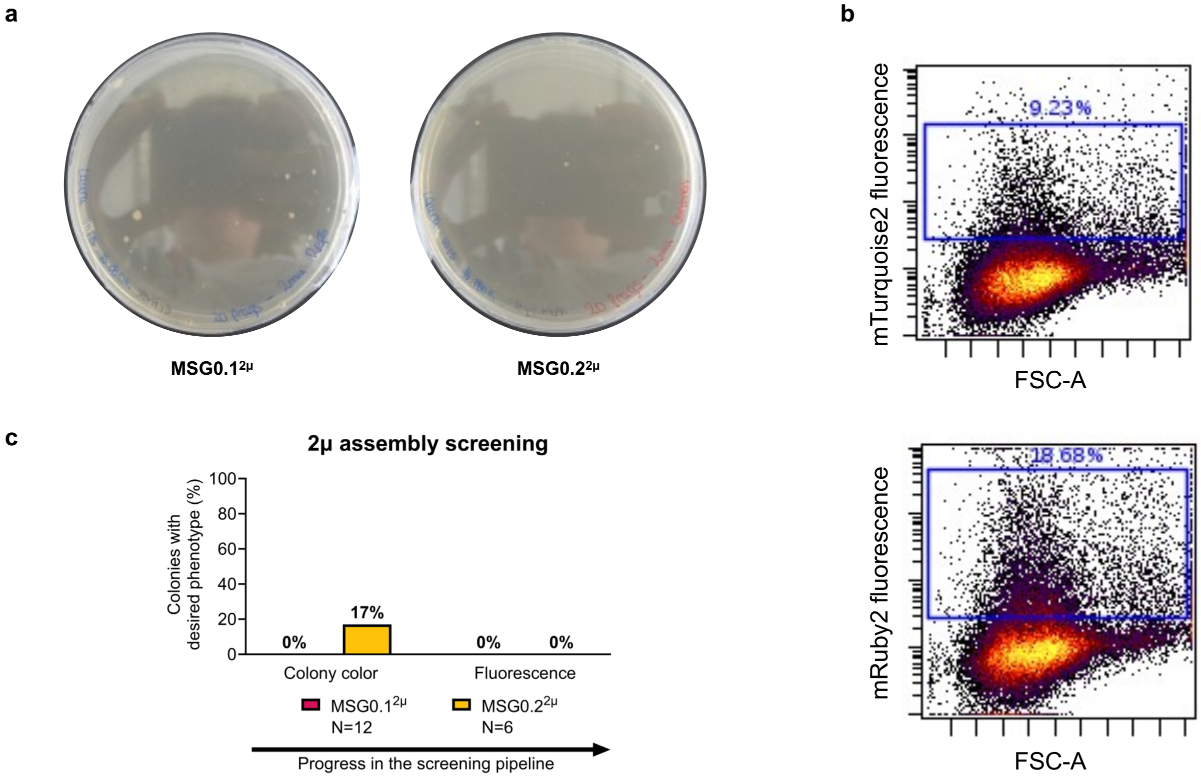


**Supplementary Figure 3: Screening for correctly assembled SynChrs with 2µ origin. a)** Plates with YPD and hygromycin after transformation. MSG0.1$^{2\mu}$ (left) and MSG0.2$^{2\mu}$ (right). **b)** FACS screening for mTurquoise2 and mRuby2 fluorescence of the only orange colony for MSG0.2$^{2\mu}$. Events within the blue frame are classified as fluorescent. **c)** Percentage of colonies with the desired phenotype at each screening step, compared to step 1. Colonies that grew on YPD with hygromycin plates were classified as positive when displaying an orange color and when showing fluorescence for both mTurquoise2 and mRuby2. Data from one transformation are displayed.

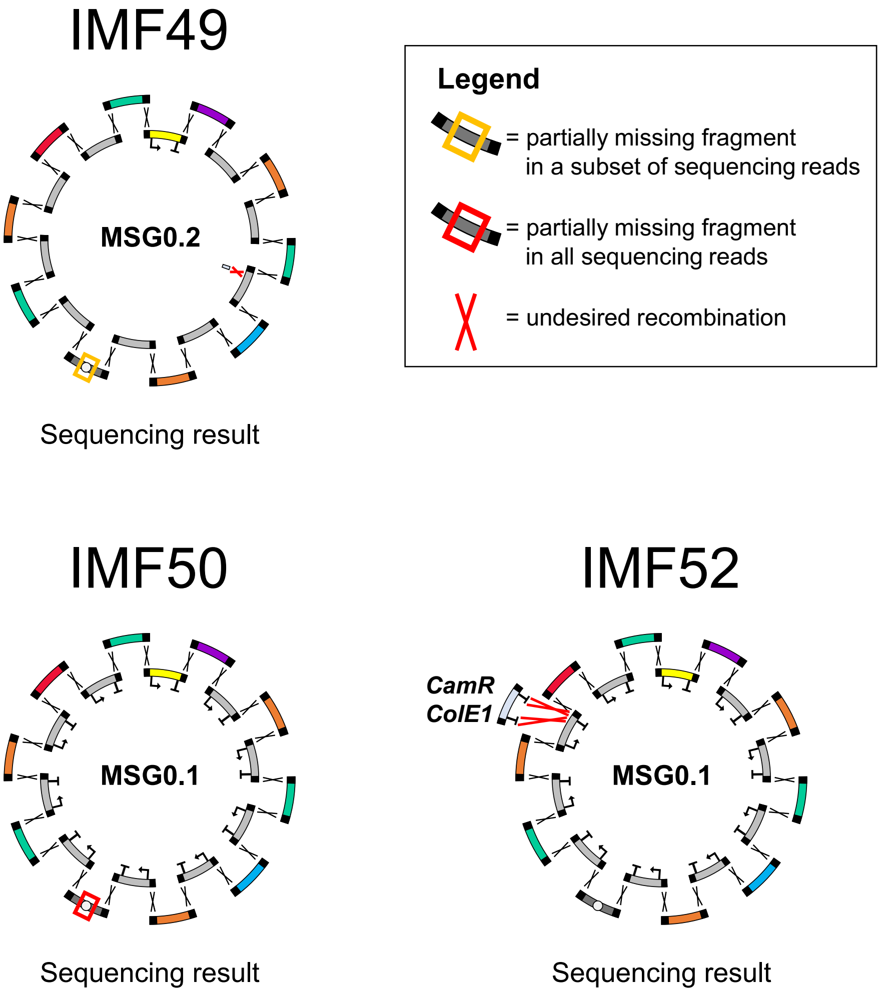


**Supplementary Figure 4: Sequenced SynChr configurations for strains IMF49, IMF50 and IMF52.** The MSG0.2 of IMF49 showed approximately half the sequencing depth for part of the CEN6/ARS4_bla_ColE1 fragment (1,891 bp) compared to the rest of the SynChr and has an 89-bp insert in CF4. MSG0.1 of IMF50 misses a 1,992-bp sequence in the middle of the fragment with CEN6/ARS4. MSG0.1 of IMF52 contains an insertion of a 1,780-bp sequence in the terminator region downstream of PF9, corresponding to a plasmid backbone (containing a chloramphenicol resistance gene and ColE1-type origin of replication), used as template during PCR.

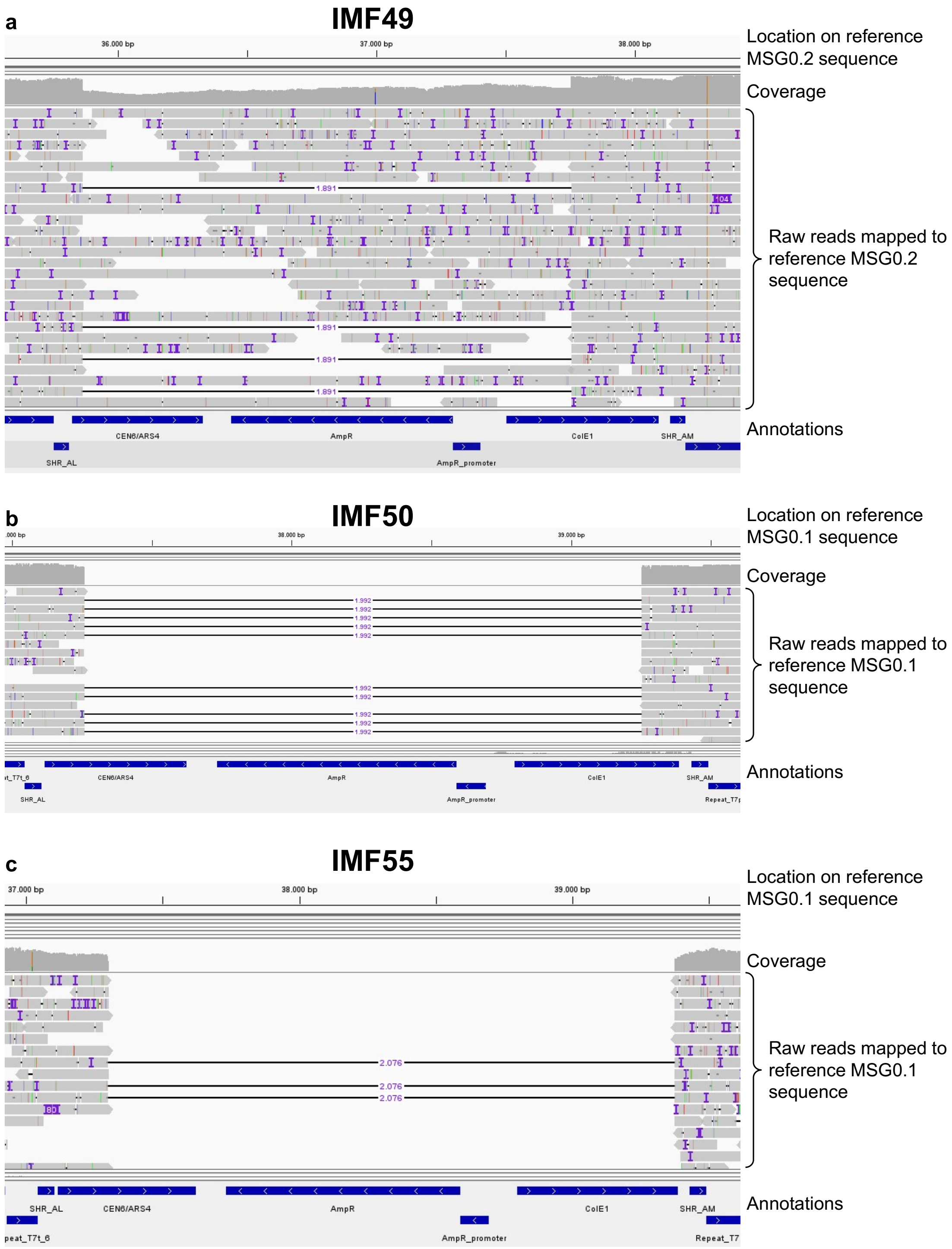


**Supplementary Figure 5: Raw reads mapped to the reference SynChr sequence, showing gaps in the CEN6/ARS4_bla_ColE1 fragment. a)** MSG0.2 from strain IMF49. A subset of the raw reads contains an 1891-bp gap from the beginning of *CEN6/ARS4* until halfway ColE1. **b)** MSG0.1 from strain IMF50. All raw reads contain a 1992-bp gap from approximately one-third of *CEN6/ARS4* until approximately three-quarters of ColE1. **c)** MSG0.1 from strain IMF55. All raw reads contain a 2076-bp gap from halfway *CEN6/ARS4* until the end of ColE1. Annotation AmpR = *bla* gene.

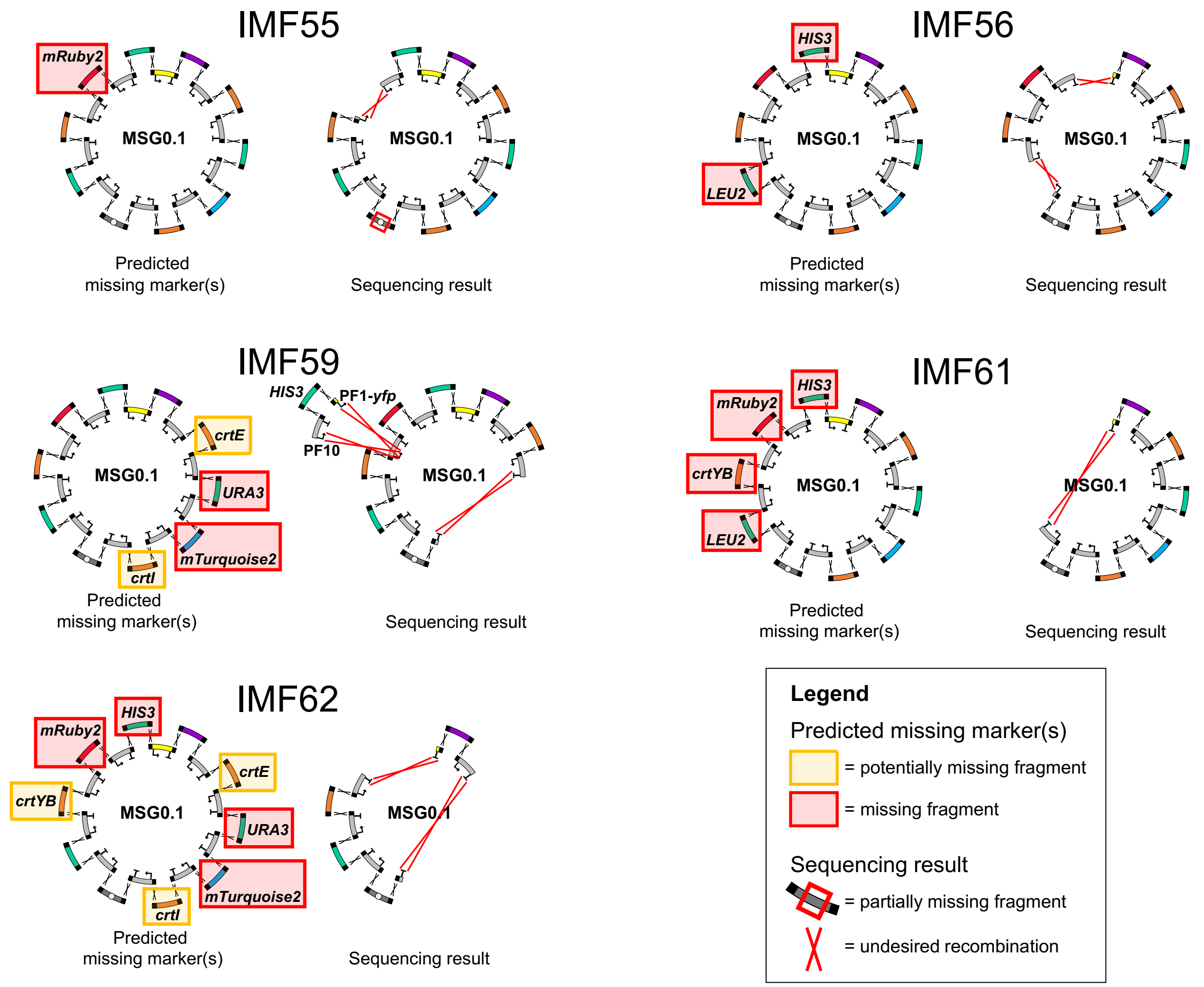


**Supplementary Figure 6: Predicted versus sequenced MSG0.1 configurations for strains missing at least one marker.** On the left the predicted missing markers. On the right the MSG0.1 configuration as revealed by sequencing.

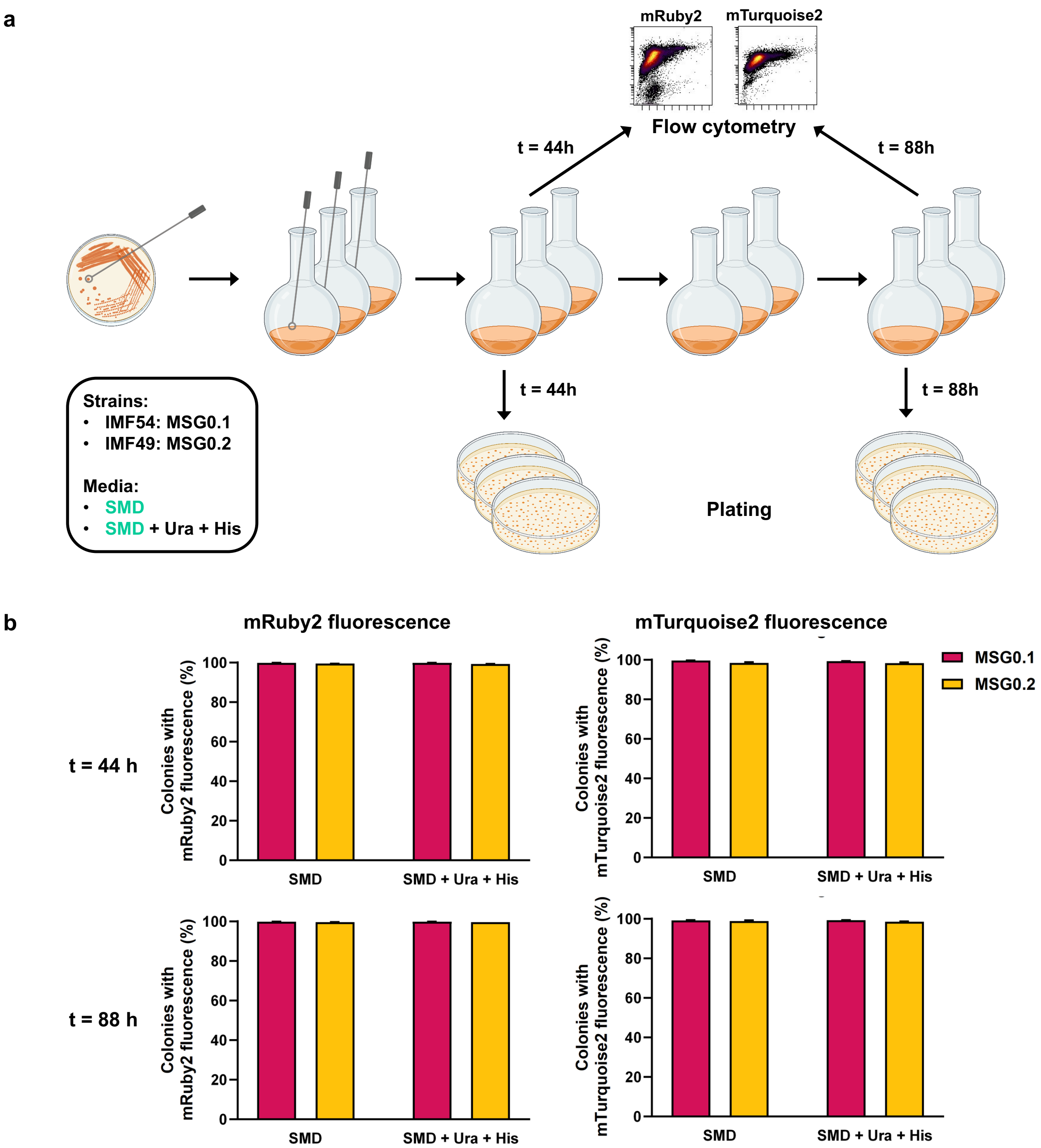


c

| | Number of white colonies after 44 h of growth | | | | Number of white colonies after 88 h of growth | | | |
|---|---|---|---|---|---|---|---|---|
| | MSG0.1 | | MSG0.2 | | MSG0.1 | | MSG0.2 | |
| | SMD | SMD + Ura + His | SMD | SMD + Ura + His | SMD | SMD + Ura + His | SMD | SMD + Ura + His |
| Plate 1 | 1 | 0 | 0 | 0 | 1 | 1 | 0 | 0 |
| Plate 2 | 0 | 1 | 0 | 0 | 0 | 0 | 0 | 0 |
| Plate 3 | 0 | 1 | 0 | 0 | 0 | 1 | 3 | 0 |

**d**

**MSG0.1, SMD, 44 h**

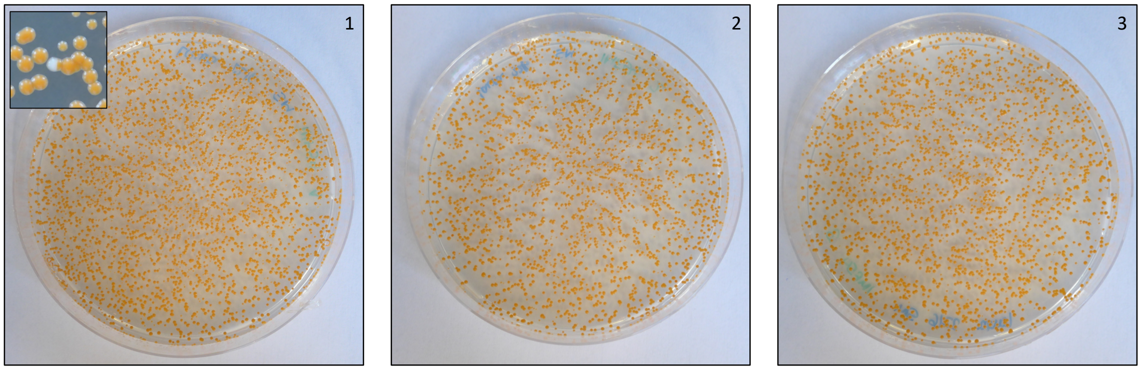


**MSG0.1, SMD + Ura + His, 44 h**

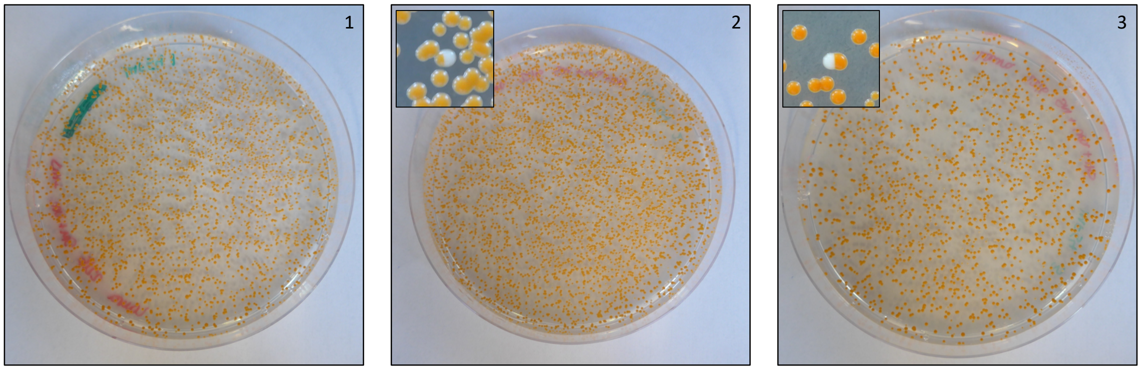


**MSG0.2, SMD, 44 h**

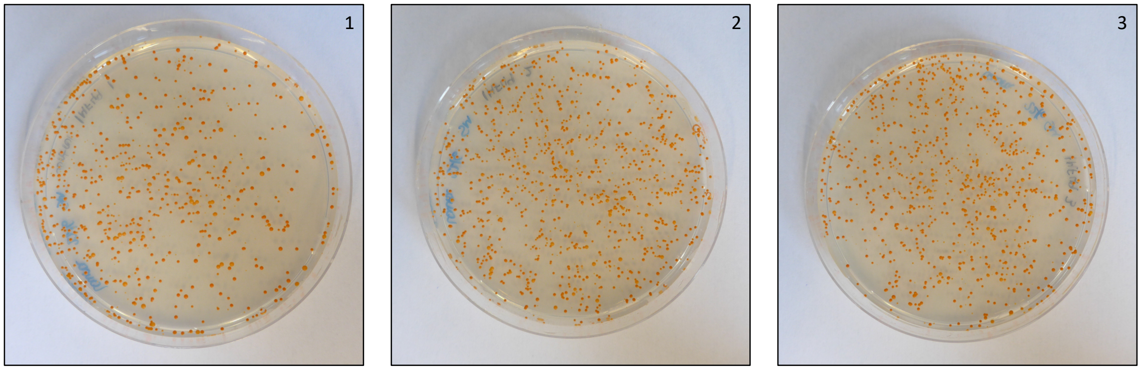


**MSG0.2, SMD + Ura + His, 44 h**

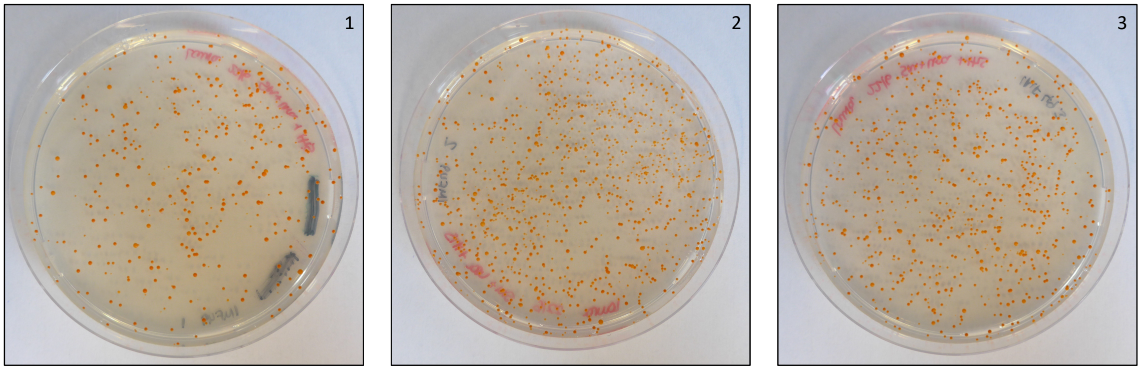

**e**

**MSG0.1, SMD, 88 h**

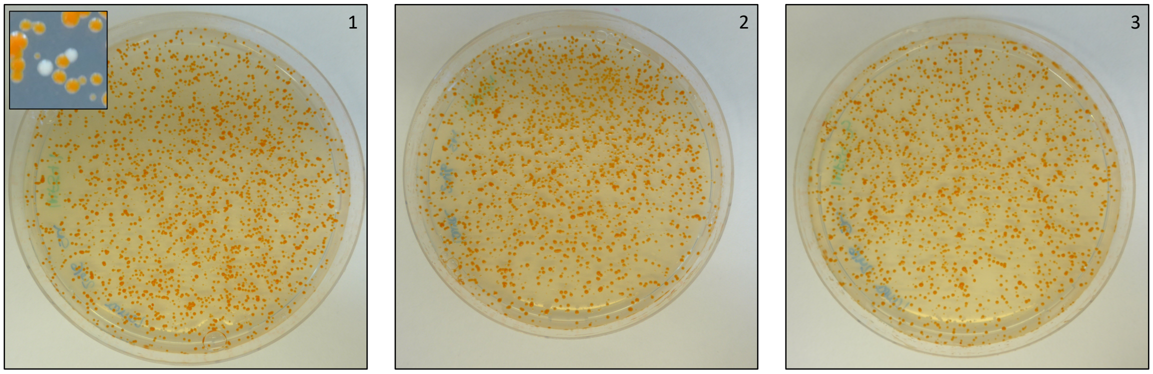


**MSG0.1, SMD + Ura + His, 88 h**

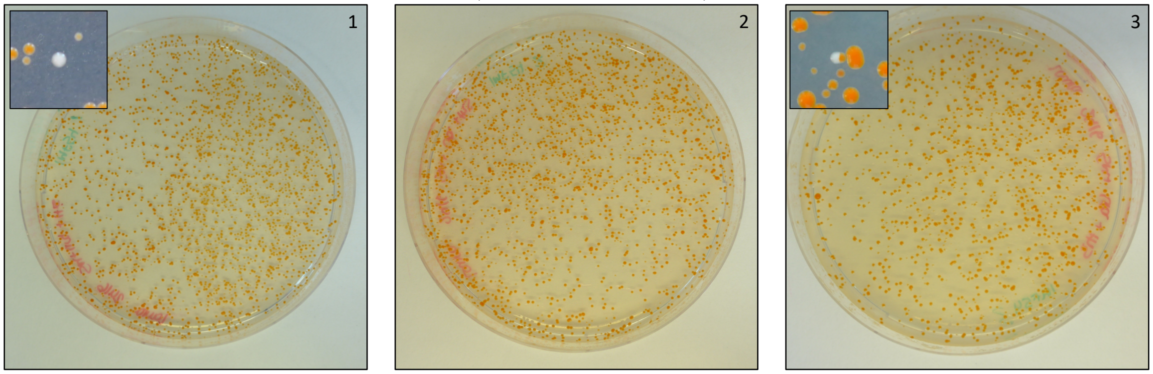


**MSG0.2, SMD, 88 h**

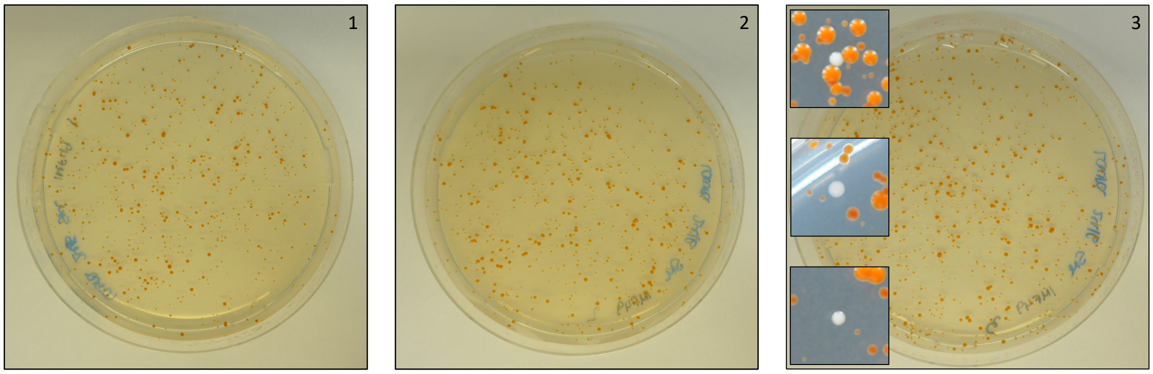


**MSG0.2, SMD + Ura + His, 88 h**

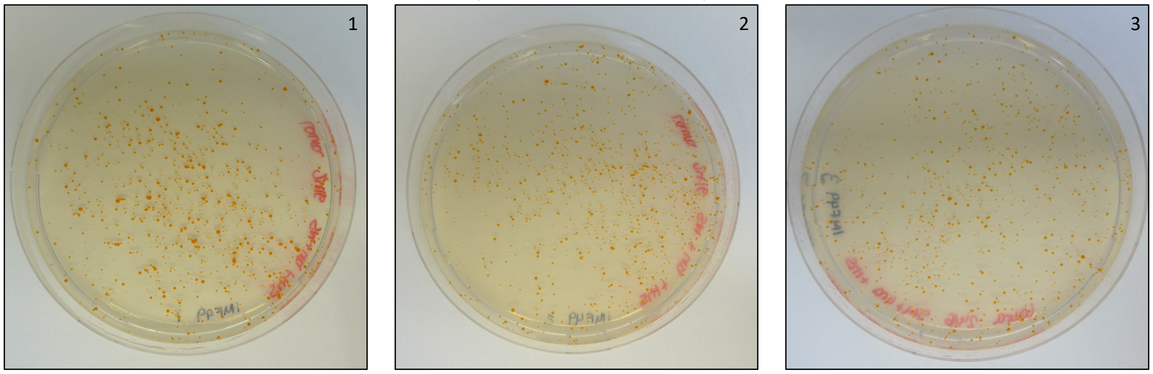

**Supplementary Figure 7: SynChrs are stable during strain propagation. a)** Experimental approach to test SynChr stability. Strains IMF54 (MSG0.1) and IMF49 (MSG0.2) were propagated for 4 days (88 h) in biological triplicates in 100 mL high-selection medium SMD and low-selection medium SMD + Ura + His. Media were refreshed every day. Samples were taken on day 2 (after 44 h) and day 4 (after 88 h), and tested for fluorescence by flow cytometry and for orange color by plating (in the same selective medium as the liquid cultures). **b)** Flow cytometry results for mRuby2 and mTurquoise2 fluorescence after 2 and 4 days (44 h and 88 h, respectively) of growth in culture, in both media conditions (SMD and SMD + Ura + His), for both strains IMF54 (SMG0.1) and IMF49 (SMG0.2). **c)** Number of white colonies per plate after 44 h or 88 h of growth in triplicates. **d) – e)** Plates showing low incidence of white colonies after 44h and 88 h of growth, respectively, in both media conditions for both strains in triplicates. Insets show only white colonies of each plate.

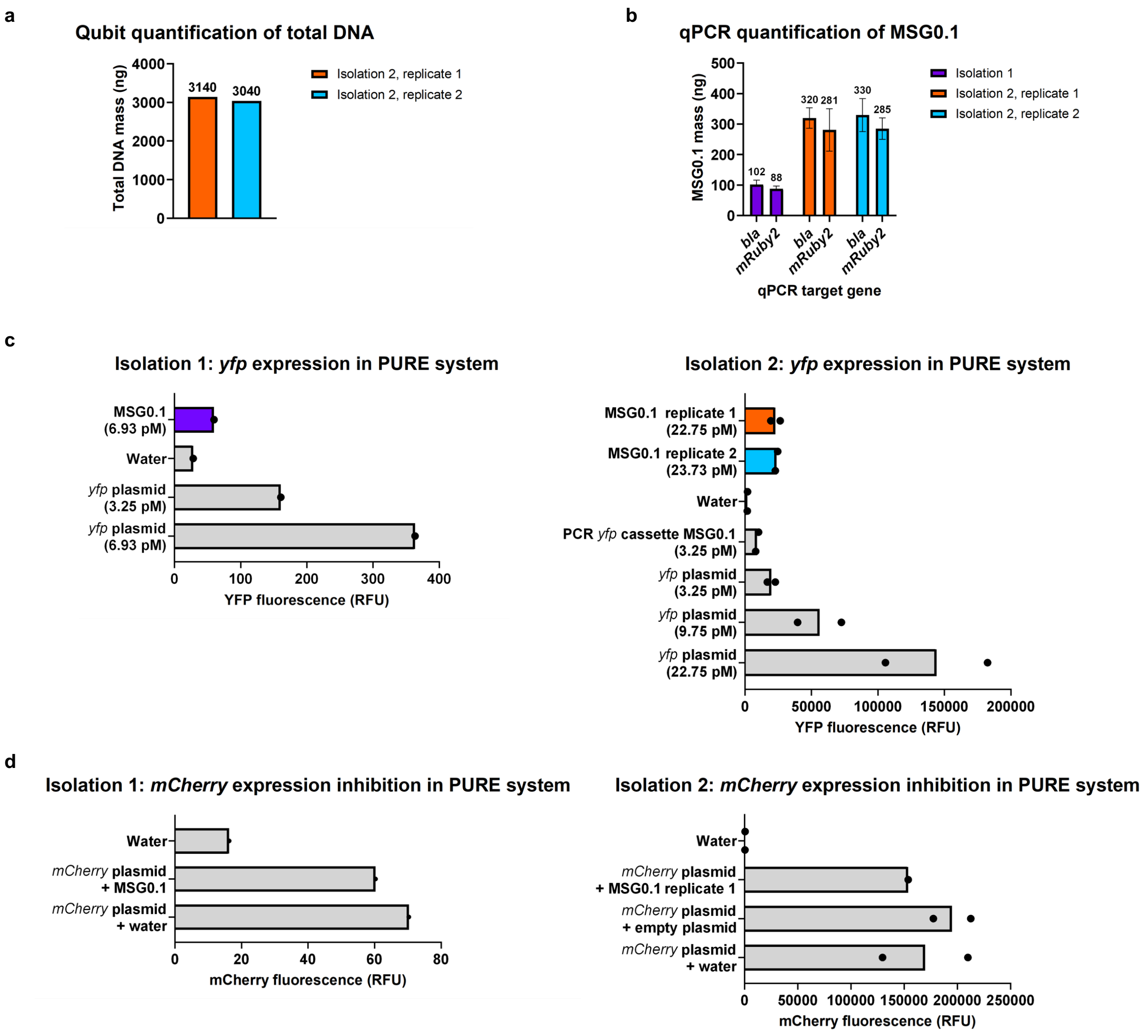


**Supplementary Figure 8: MSG0.1 can be isolated from *S. cerevisiae* and shows expression of *yfp* in PURE system.** Data from two independent isolations are presented. **a)** Qubit quantification of total isolated DNA mass (ng) in 100 µL for isolation 2. Data from two biological replicates is shown. **b)** qPCR quantification of MSG0.1 mass (ng) in 100 µL, quantified by primer sets targeting *bla* and mRuby2 genes. Mean and standard deviation from three technical replicates are plotted. **c)** YFP fluorescence measured after 16h incubation of various DNA templates in PURE system. Individual data points indicate independent PURE reactions. Note that different plate readers were used for isolation 1 and 2. Final DNA template concentrations in the assembled PURE reaction are indicated between brackets. MSG0.1 = MSG0.1 is isolated from *S. cerevisiae* strain IMF54. Water = No DNA template (negative control). *yfp* plasmid = Plasmid containing a *yfp* gene, isolated from *E. coli* (positive control). PCR *yfp* cassette MSG0.1 = *yfp* cassette amplified by PCR from MSG0.1 isolated from *S. cerevisiae* strain IMF54 (positive control). **d)** mCherry fluorescence measured after 16h incubation of a diluted mCherry plasmid in PURE system. Individual data points indicate independent PURE reactions. Water = No DNA template (negative control). *mCherry* plasmid + MSG0.1 = 0.5 µL of *mCherry* plasmid (final concentration: 1 nM) isolated from *E. coli*, diluted in 2.75 µL MSG0.1 sample. *mCherry* plasmid + water = 0.5 µL of *mCherry* plasmid (final concentration: 1 nM) isolated from *E. coli*, diluted in 2.75 µL water (positive control). *mCherry* plasmid + empty plasmid = 0.5 µL of *mCherry* plasmid (final concentration: 1 nM) isolated from *E. coli*, diluted in 2.75 µL non-coding plasmid.

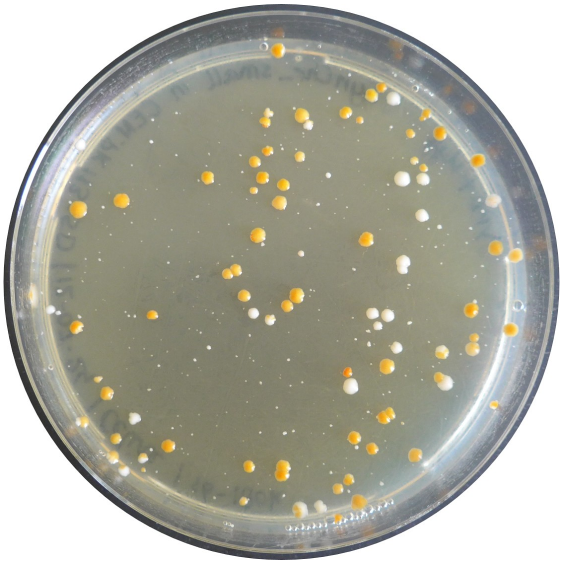

**Supplementary Figure 9: Plate after yeast transformation with assembly fragments for MSG1.**

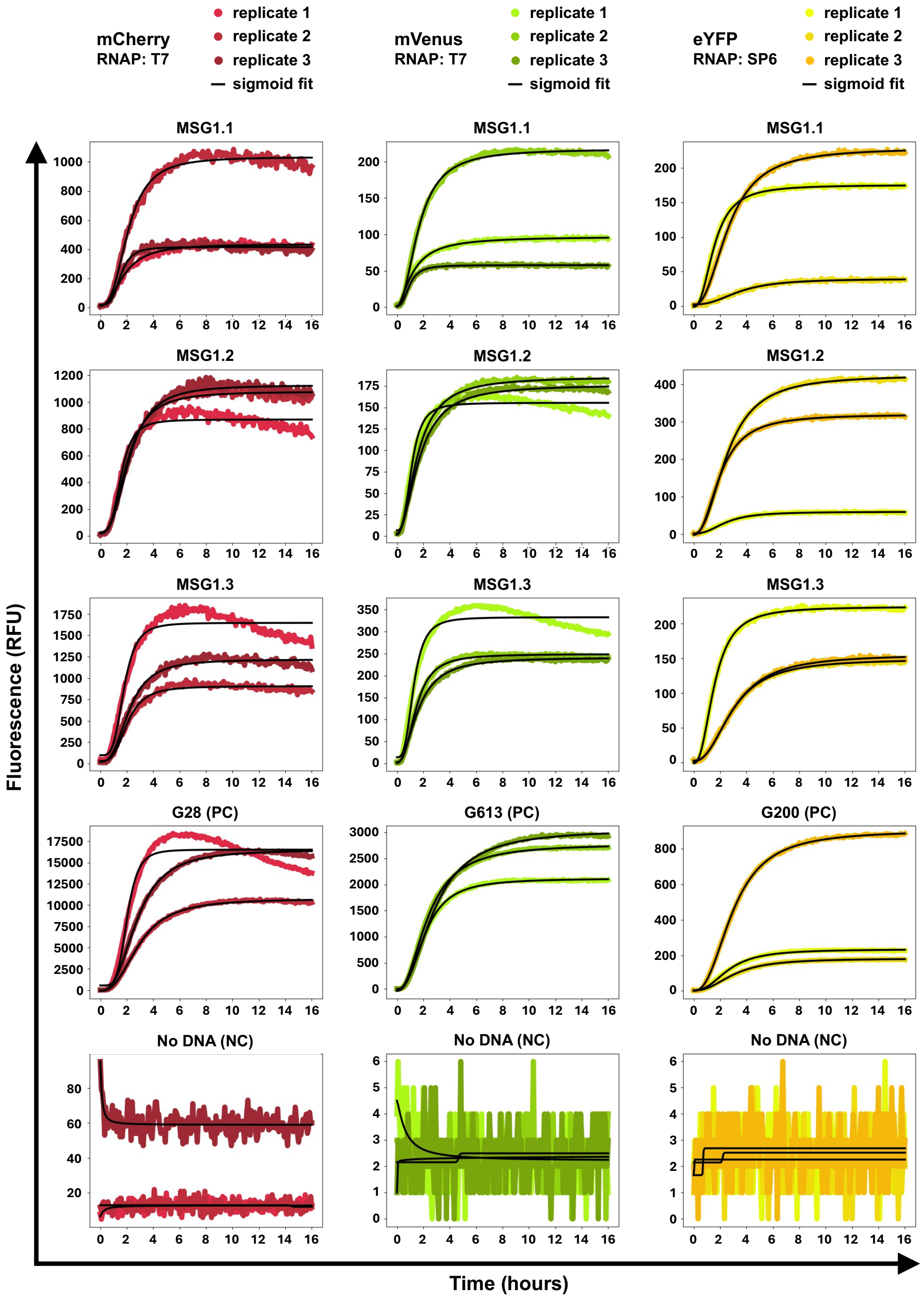


**Supplementary Figure 10: Fluorescence measurements over 16 h of expression with PURE system in bulk reactions.** The DNA templates (MSG1 variants or control DNA templates at 1 nM final concentration), as well as the fluorescent reporter proteins, are indicated. Black lines show sigmoid fitting for calculation of the apparent translation rate (maximum slope). Minimum and maximum of y-axis differ per graph for better visualization of the fluorescence profiles. Three independent PURE reactions were performed for each DNA template.

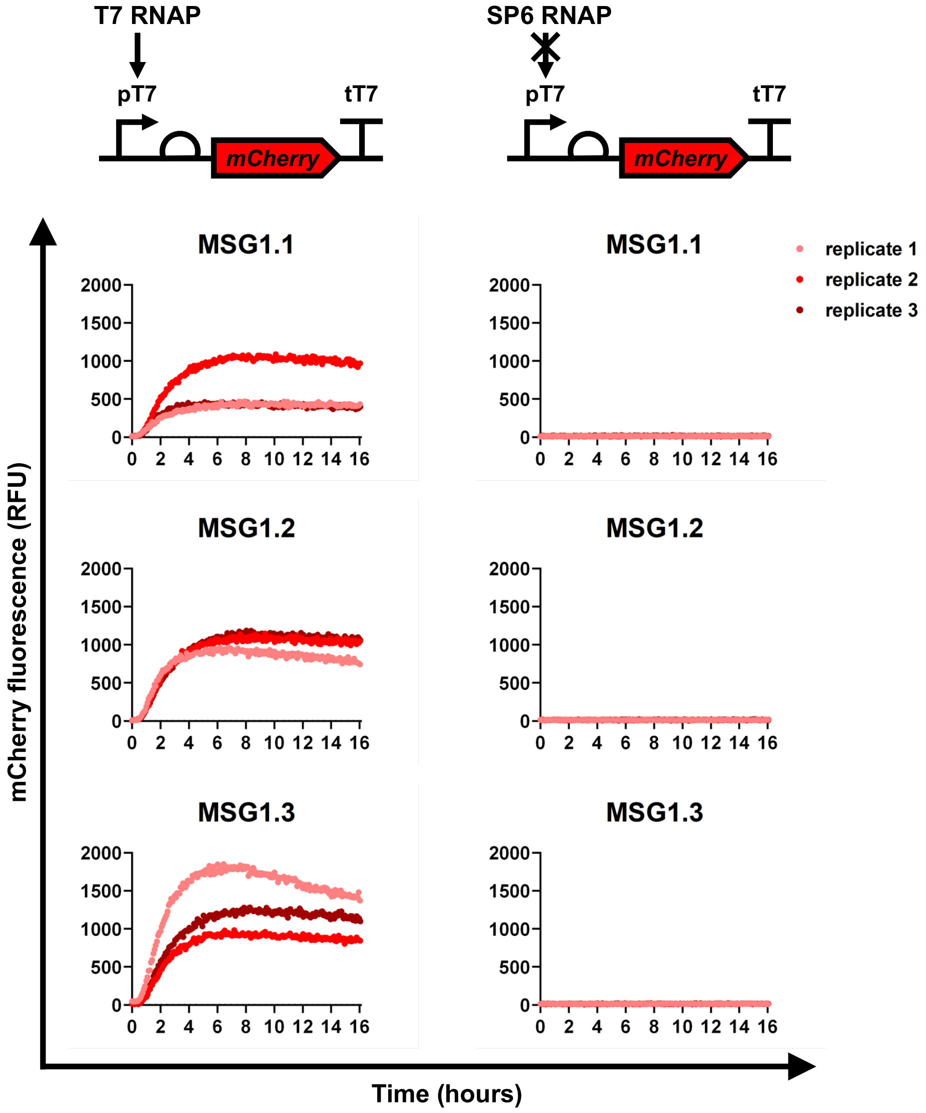


**Supplementary Figure 11: Orthogonality of SP6 RNAP with the T7 promoter.** For comparison, y-axis minimum and maximum values are the same for all graphs.

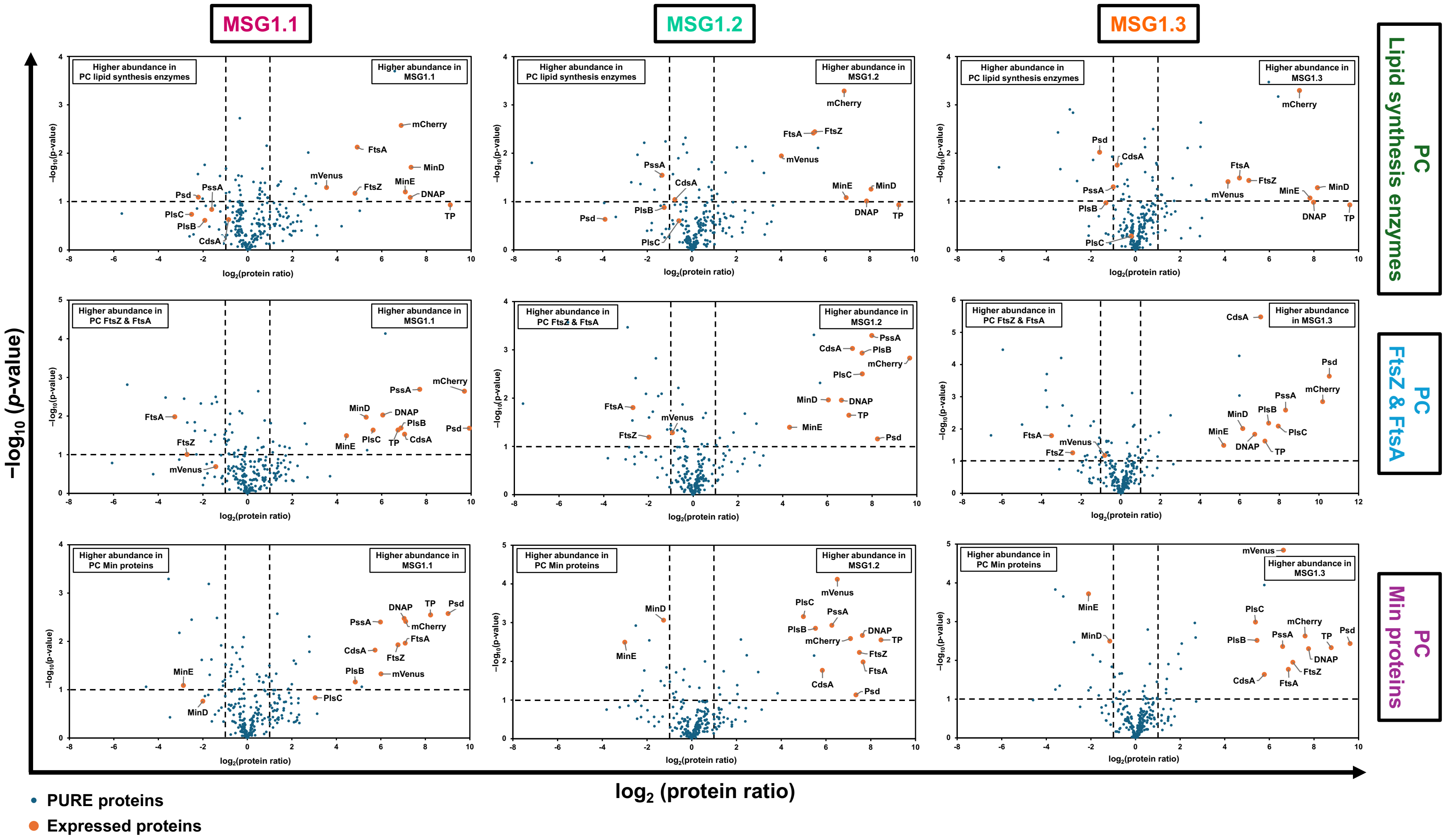


**Supplementary Figure 12: Volcano plots displaying the change in relative protein abundance between MSG1 variants and control plasmids expressed with T7 RNAP in PURE reactions after 16 h incubation.** Vertical dashed lines indicate a 2-fold increase. The horizontal dashed line indicates a *p*-value = 0.1 from a two-tailed *t*-test. Values are calculated from independent PURE reactions ($n$). $n$ = 2 for MSG1.1 and the control plasmid for lipid synthesis enzymes, and $n$ = 3 for all other conditions. Because $n$ = 1 for the control plasmid for DNA replication proteins, no volcano plot could be generated for this condition. PC, positive control

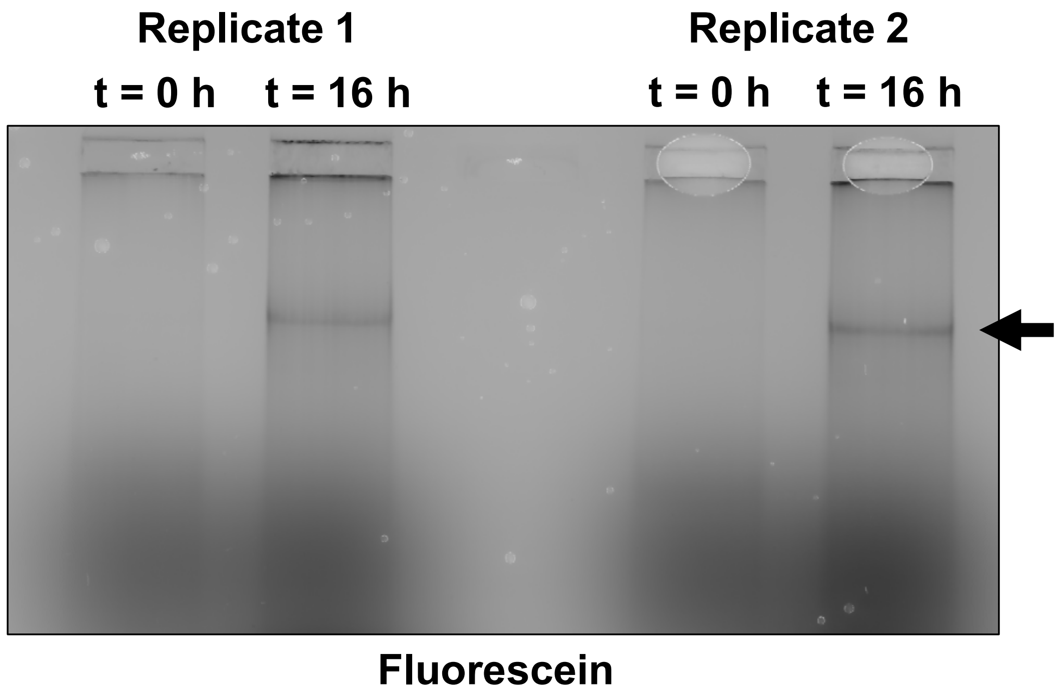


**Supplementary Figure 13: Full-length synthesis of MSG1.1 visualized on agarose gel.** The fluorescein channel shows newly synthesized DNA. The arrow points at the expected band for linear MSG1.1 with a size of 41 kb. The reaction contained ribosomes and was performed in duplicate.
